# Sequencing-free Tissue-wide Spatial Profiling of Post-transcriptional Regulations

**DOI:** 10.1101/2022.12.11.519542

**Authors:** Xianglin Ji, Peilin Fang, Xi Zhao, Chuanyin Xiong, Qi Yang, Youyang Wan, Richard Yan Do, Zixun Wang, Lin Qi, Linfeng Huang, Wenjun Zhang, Xin Wang, Peng Shi

## Abstract

The importance of genetic or epi-genetic heterogeneity has been increasingly recognized, but it has been challenging to profile intracellular post-transcriptional targets with sufficient throughput and resolution at across large-scale tissue samples. This study describes a technique, Spectrum-FISH, for high-throughput, sequencing-free, and tissue-wide spatial profiling of various post-transcriptional targets in acute tissue sections with subcellular resolution. The platform uses a biochip with an array of vertically aligned nanoprobes to effectively extract intracellular molecules for downstream analysis in the coordinates of the large-scale of cells within a tissue slice. As a proof-of-concept, the Spectrum-FISH is used to profile the spatial dynamics of 24 miRNAs and 9 m^6^A-modified messenger RNAs (m^6^A-mRNA) in acute olfactory bulb (OB) slices of millimeter scale. The results showed potentially multiomics spatial heterogeneity for the examined post-transcriptional regulations in rodent OB, especially in the outer plexiform layer and granule layer, where highly correlated miRNAs and m^6^A-mRNAs groups were identified, indicating a potential cooperative involvement of different post-transcriptional regulations at these OB regions.

## Introduction

Spatial heterogeneity in gene expression is closely related to human physiology in normal or diseased conditions ^1^. The development of techniques for capturing such spatial information with cellular resolution across tissues becomes increasingly more important for advancing our knowledge regarding critical genetic or epigenetic regulations among large-scale cellular populations. Particularly, fluorescence in situ hybridization (FISH) has been widely used to assess the spatial distribution of mRNAs ^2–4^, further developments of FISH with different encoding strategies have enabled highly multiplexed analysis of mRNAs ^5, 6^. MERFISH (multiplexed error-robust FISH) uses combinatorial labeling with an error-robust algorithm for simultaneous imaging RNA species in a single cell ^5^. SeqFISH (sequential fluorescence in situ hybridization) is capable of sequential analysis of different RNAs through multiple rounds of hybridization to increase the assay throughput ^6^. FISH-based profiling technologies mostly focus on the subcellular localization of genetic targets and are applied to individual cells, which is not ideal for profiling the heterogeneity of wide tissue samples. In some cases^5–7^, the requirement of high-sensitivity single-molecule fluorescence imaging systems (e.g. super resolution microscopy) poses further technical barriers for its wider applications^8^.

On a larger scale, tissue-wide spatial transcriptomics has been recently developed ^8–14^. A typical implementation uses arrayed poly(T) tails to capture released mRNAs from a pre-treated tissue sample on a grid of spatially-indexed coordinates, and the captured mRNAs are later identified by sequencing ^9, 10^. While useful, the technique requires complex histology processing of tissue samples, involving frozen sectioning, fixation, and membrane penetration, to release intracellular mRNAs. Not only laborious and time-consuming, these procedures may also lead to cross-contamination among adjacent regions or RNA degradation over certain spatial extent, thus reducing the spatial resolution of associated transcriptomic analysis ^9, 15^. Moreover, the majority of tissue-wide spatial profiling techniques are limited to transcriptional (mRNAs), proteomic targets ^8, 16^ or chromatin modification^17^, and it is still challenging to analyze post-transcriptional regulations at a similar scale.

Out of various post-transcriptional regulations, microRNAs (miRNAs) and RNA methylation are prevalent mechanisms that are closely related to the generation of complex gene expression topology to accommodate substantial regulatory flexibility, diversity, and robustness ^18^. miRNAs are noncoding single-stranded small RNA molecules (∼21–23 nucleotides) that modulate gene expression by inhibiting mRNA translation ^19^. RNA methylation is the most common RNA modification on mRNAs for regulating their metabolism. For example, N^6^-methyladenosine (m^6^A) modification is highly enriched in the mammalian brain, and distinct m^6^A methylation patterns varies at different brain regions with a dynamic involvement in neural development ^20^. Recently, the cooperative involvement of the two mechanisms has also been identified to play important roles in post-transcriptional genetic regulation^18^. However, for either miRNAs or m^6^A methylation, there are very limited techniques that can analyze the associated post-transcriptional regulation with sufficient throughput and spatial information at a tissue-wide scale ^15, 21^, and even less tools are available for correlated spatial analysis of multiple post-transcriptional regulations. Particularly, the requirement for long priming probes in FISH-based methods prevents its application towards small miRNAs. For mRNAs with m^6^A modifications, although the nucleotide length is sufficient for probe priming in FISH, additional biochemical analysis, such as liquid chromatography, need to differentiate the methylation levels. The recently developed spatial transcriptomic techniques are not yet applicable to profiling the subtle details of post-transcriptional processes regulated by miRNAs or methylated modifications ^8–10^.

Here, we present a general platform, Spectrum-FISH (digital spectrum barcoded molecular fishing), for large-scale profiling of post-transcriptional regulations with subcellular spatial resolution across centimeter-wide fresh tissue samples. The platform is based on a molecular fishing system using an array of vertically-aligned nanoprobes ^22–24^, which are capable of high-throughput isolation of intracellular molecules by interfacing with the superficial cell layer in acute tissues slices. With different functionalization, the molecular fishing is tailored to capture miRNAs or m^6^A modified mRNAs, which are in situ analyzed on the nanoprobes. The resulted molecular analysis is then registered with subcellular spatial resolution to a large population of individual cells, providing a tissue-wide spatial epi-transcriptomics of the associated molecular targets. For multiplexing, a spectrum barcoding system enables digitized quantification of the molecular targets on each nanoprobes, which are individually mapped to single cells by associative reference to a spatial coordinate established by imprinting the tissue geometric features on the nanoprobe array. In this proof-of-concept study, the Spectrum-FISH technique is used to profile the spatial expression of multiple miRNAs and m^6^A-modified messenger RNAs (m^6^A-mRNA) with single cell resolution in acute olfactory bulb (OB) slices of millimeter scales. The results showed potentially multiomics spatial heterogeneity for the examined post-transcriptional regulations in wild type rodent OB, especially in the outer plexiform layer (OPL) and granule layer (GR), where highly correlated miRNAs and m^6^A-mRNAs groups were identified, indicating a potential cooperative involvement of different but correlated post-transcriptional regulations at particular OB regions. This study is a first demonstration of tissue-wide spatial profiling of post-transcriptional regulations. With little adaption, the Spectrum-FISH technique can be readily extended to analyze a wide range of biological targets, to provide new insights towards their single or combinatory roles in normal or diseased conditions.

## Results

### Biochips with vertically aligned nanoprobes for post-transcriptional profiling

The Spectrum-FISH technique for post-transcriptional profiling was extended from the “molecular fishing” platform, where a random array of vertically aligned diamond nanoprobes was used to capture targeted biomarker molecules from living cells in a minimum invasive format ^22–24^. In this study, an orderly array of silicon nanoprobes was utilized to disrupt cell membrane for access to intracellular domain and to extract targets of interest from individual cells (**Figure 1a, b, Supplementary Figure S1, Supplementary Note 1**). Specifically for acute tissue samples, all the cells under the coverage of the biochip are simultaneously examined with a single operation. Unlike existing spatial transcriptomics technologies, where the tissue samples typically require extensive pre-treatment before genetic or transcriptional analysis ^8–10^, our Spectrum-FISH directly work on freshly prepared acute tissue slices. The nanoprobes were fabricated as a square array of vertical cylinders (∼800 nm diameter, ∼7 μm height, 5 μm spacing, **Supplementary Figure S2**). Depending on the targets to be analyzed, different ‘bait’ molecules are functionalized on the nanoprobes for molecular extraction. Here, p19 protein was used to isolate miRNAs and specific antibodies were used to isolate RNAs with m^6^A modifications.

**Figure 1.**
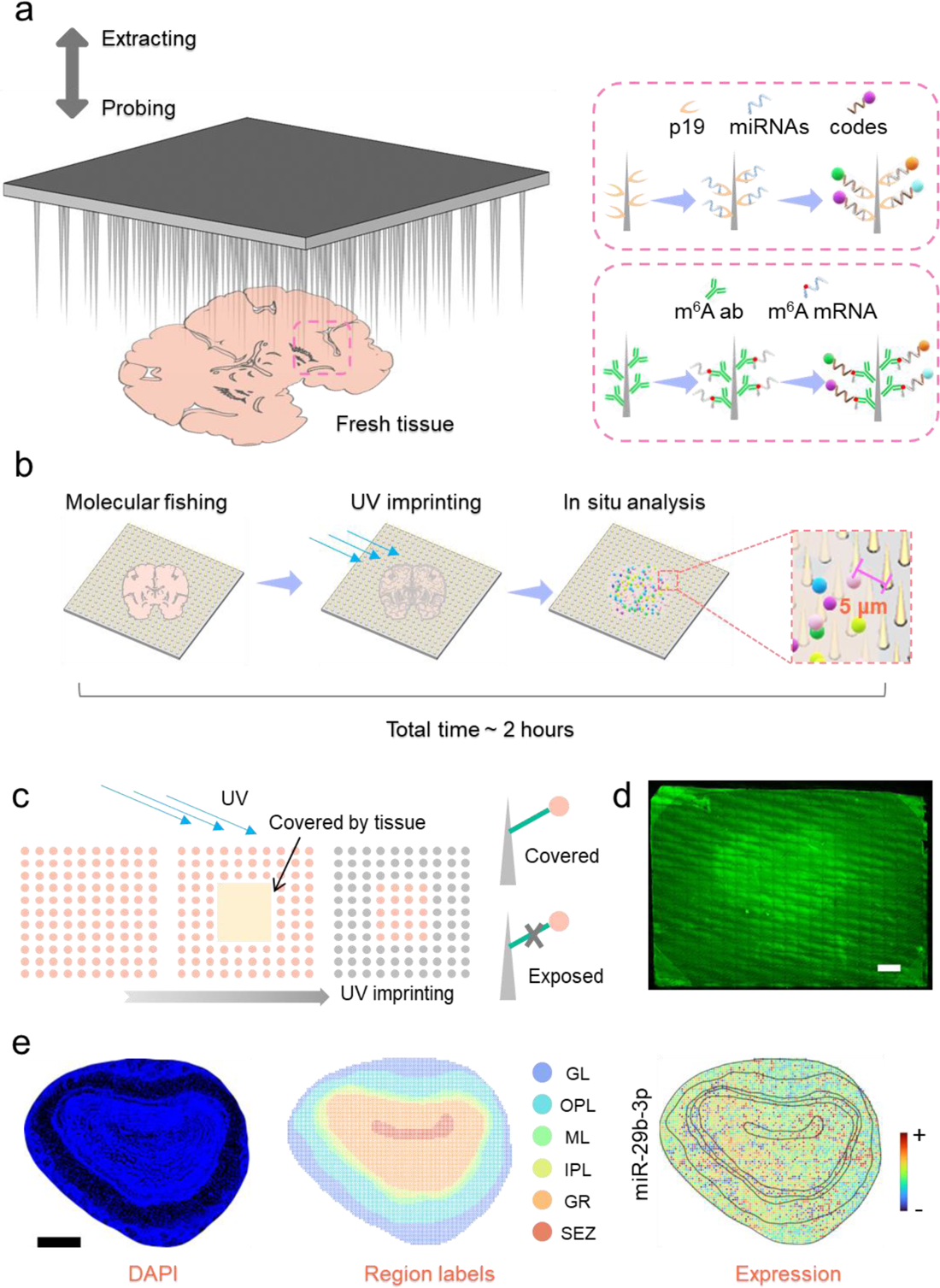
Overview of the Spectrum-FISH technique for tissue-wide spatial profiling of post-transcriptional regulations. **(a)** Procedure schematics for the molecular fishing based intracellular extraction of miRNAs or m^6^A-mRNAs from acute tissue slices. An array of vertical silicon nanoprobes were functionalized with ‘bait’ (RNA binding protein, p19 or m^6^A antibody) to directly pull multiple targeted miRNAs or m^6^A mRNAs out of cell cytoplasm in a few minutes and finally quantified by Spectrum-CODEs. **(b)** Workflow of a Spectrum-FISH analysis consist of molecular fishing, UV imprinting and in situ spatial and expressional analysis. “molecular fishing” system uses an array of silicon nanoprobes as fishing rods for minimum-invasive access of cytoplasmic regions of mammalian cells. Specifically, for “fishing” miRNAs, a size-dependent RNA binding protein, p19, was cross-linked to functionalize the nanoprobes, working as the “bait” to capture double-strand RNAs (dsRNAs). In the Spectrum-FISH system, just after the molecular fishing procedure when the tissue was still in contact with the biochip, the spatial encoding was facilitated by a UV imprinting process, which gives a snapshot of the tissue morphology and boundary on the nanoprobe biochip and provides an associative reference linking the cell and nanoprobe coordinate. After the UV imprinting procedure, the silicon nanoprobes biochip was separated from the tissue slice with the target molecules extracted by specifically binding with the ‘bait’. For multiplex visualization and quantification of the captured biotargets, we developed a digital spectrum method (Spectrum-CODE) for functional encoding the analytical targets. **(c)** Schematic illustration of the UV imprinting facilitated spatial encoding. The nanoprobes were labelled with fluorophores by a photocleavable crosslinker. When a piece of tissue slice (smaller than the chip) was interfaced with the nanoprobes for “molecular fishing”, a brief UV (365 nm, ∼5 mw/cm^2^) irradiation was applied. This energy level was just enough to cleave off the fluorescent labels on the nanoprobes in direct UV exposure, while the rest part covered by the tissue remained unaffected, generating a pixelated fluorescent pattern of the tissue, which can be used to register of the nanoprobes to individual cells at later analytical stages. **(d)** Representative fluorescence images showing the pixelated morphology and boundary features of a tissue slice on the nanoprobe biochip. Scalebar, 400 μm. **(e)** Example results from miRNA spatial profiling in an acute OB slice. Left: nuclei staining by DAPI; middle: OB region labels; right: expression of miR-29b-3p across an OB slide. Scalebar, 400 μm.

For spatial epi-transcriptome analysis at single cell level, the key is to map the nanoprobes in the coordinates associated with the large number of cells with microscale resolution. In the Spectrum-FISH system, the spatial encoding was facilitated by a UV imprinting process, which gives a snapshot of the tissue morphology and boundary on the nanoprobe biochip and provides an associative reference linking the cell and nanoprobe spatial information. The nanoprobes were labelled with fluorophores (Alexa Fluor 568, AF-568) by a photocleavable crosslinker (**Figure 1b, c**). When a piece of tissue slice (smaller than the chip) was interfaced with the nanoprobes for “molecular fishing”, a brief UV (365 nm, ^∼^5 mw/cm^2^) irradiation was applied. This energy level was just enough to cleave off the fluorescent labels on the nanoprobes in direct UV exposure, while the rest part covered by the tissue remained unaffected (**Figure 1c**), generating a pixelated fluorescent pattern of the tissue (**Figure 1d, Supplementary Figure S3**), which can be used to register of the nanoprobes to individual cells at later analytical stages. In this way, the nanoprobe-associated post-transcriptional profiles can be spatially decoded at a single cell level (**Figure 1b, e**), as shown in a representative analysis of miRNA (miR-29b-3p) expression in an OB acute slice.

### High-throughput multiplexing based on spectrum digitization

After a “molecular fishing” operation, the biochemical signals captured on the nanoprobes were visualized and quantified by fluorescence imaging. To overcome the challenge of limited numbers of fluorescent channels and increase the multiplexing throughput of the Spectrum-FISH technique, we developed a digital spectrum method for functional encoding the analytical targets (**Supplementary Note 2**). The extracted biomarker molecules (e.g. miRNA or m^6^A-mRNA) were visualized by an in-situ hybridization with a “reporter probe” DNA, which was specifically designed to recognize and encode different targets. Also, each probe sequence was pre-labelled by amino-Fe_3_O_4_ nanobeads with a rainbow fluorescence composed by a mixed ratio of the 4 different fluorophores, including AF-488, 514, 555 and 647 (**Figure 2a, b, Supplementary Figure S4**). In the simplest case, the digitized spectrum uses a “1” or “0” encoding strategy to indicate the presence of a particular fluorophore in the labeling process (**Figure 2c**). Particularly for analyzing RNAs, the DNA probes resulted from one round of hybridization can later be removed by a DNase treatment to further expand of the analytical throughput by multiple rounds of probe hybridization towards more targets (**Supplementary Figure S5**). Theoretically, a multi-channel system could render the number of codes totaled *N* × (2^C^ − 1), where *C* is the number of fluorescent channels, *N* is the rounds of the hybridization analysis, and code “0000” (blank, no fluorescence) is not used. In the four-channel system used in this study, the rainbow fluorescent beads were encoded by 15 spectral combinations for analysis (**Figure 2d**). In the binary encoded labelling, the fluorescence signal from each bead was respectively acquired from 4 spectral windows, each of which worked as a digital input of a 4-dimensional spectral vector (**Figure 2c, d, Supplementary Figure S4**). With our custom developed computational analysis, the digitized spectral vectors of all the beads can be well separated (**Figure 2e**). To overcome the interference by the variation of fluorescence intensities and a low level of background signal, especially in the “0” coded spectral window, we used a machine learning based algorithm to differentiate the digital spectral features (**Figure 2f, Supplementary Figure S6, S7**), which provided autonomous recognition of the digital spectrum codes with nearly 90% accuracy. For a small fraction of uncertain cases, a ‘filtering’ step (**Supplementary Figure S8, S9**) was further used to remove the beads with ambiguous signal that did not carry a typical spectral feature or simply the noise resulted from microscopic imaging (**Supplementary Figure S10**). After spectrum decoding, for each target, the number of nanobeads on individual nanoprobes were counted to indicate the copy number of the molecular targets (e.g. miRNA or mRNA).

**Figure 2.**
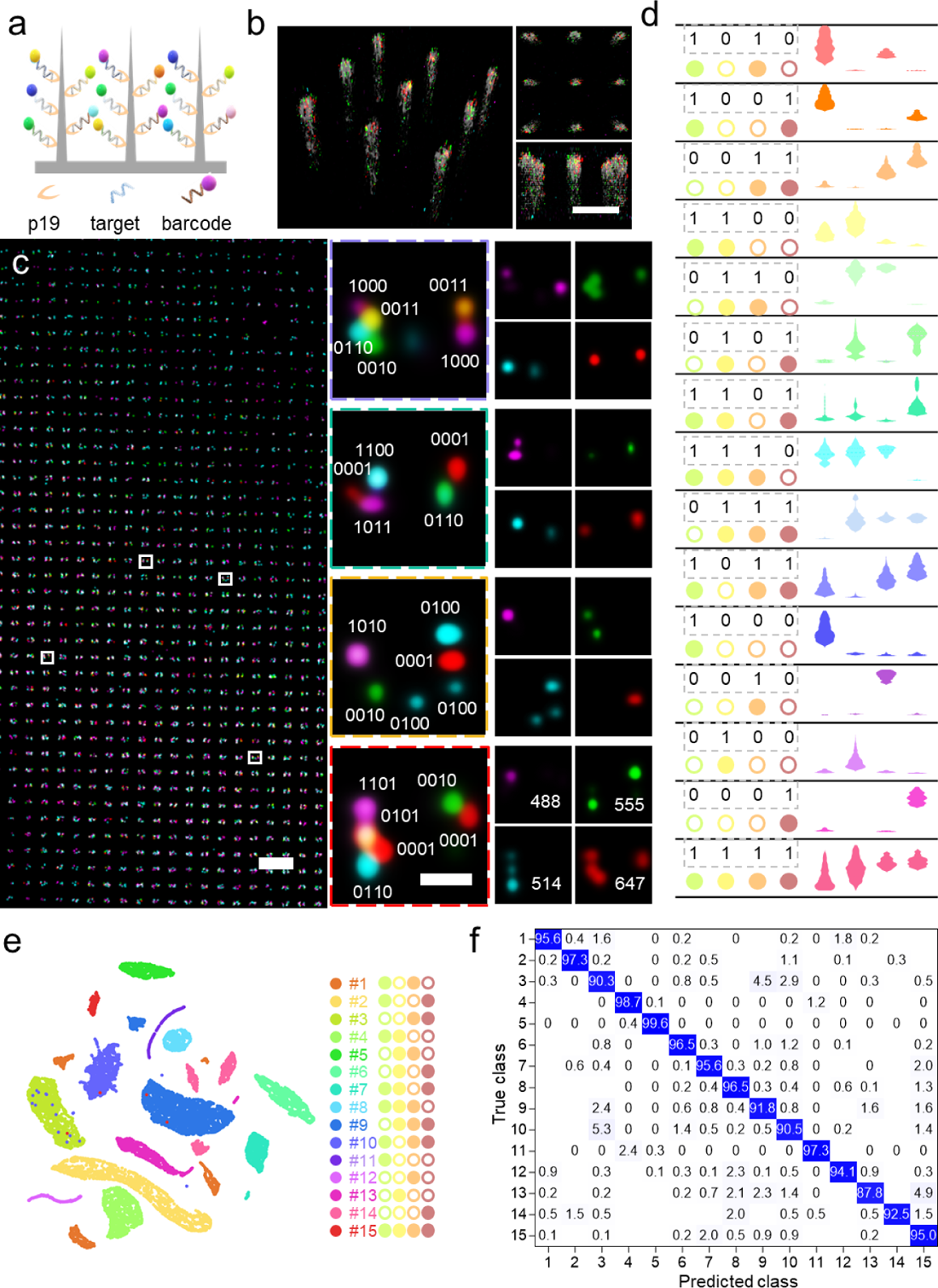
High-throughput multiplexing based on spectrum digitization. **(a)** The multiplexing strategy (Spectrum-CODE) using nanobeads with rainbow fluorescence labeling to encode different molecular targets. The Spectrum-CODE can overcome the challenge of limited numbers of fluorescent channels and increase the multiplexing throughput of the Spectrum-FISH technique, The extracted biomarker molecules (e.g. miRNA or m^6^A-mRNA) were visualized by an in-situ hybridization with a “reporter” DNA specifically designed to recognize and encode different targets. Each ‘reporter’ sequence was pre-labelled by amino-Fe_3_O_4_ nanobeads with a rainbow fluorescence composition. **(b)** Three-dimensional visualization of the spectrum encoded nanobeads after being captured and imaged on the nanoprobes. Scale bar, 5 μm. **(c)** Large scale representative fluorescence image showing the nanoprobes with multiple nanobeads encoded with different digital spectrums composed of four fluorescence channels, including Alexa fluor 488, 514, 555 and 647. Scalebar, 10 μm (left). The boxed region in the left panel is enlarged in the middle column (scalebar, 500 nm), and the right panels show the split individual fluorescent channels. **(d)** Violin plots showing the fluorescence vectors and associated barcodes for 15 types of nanobeads with different digital spectrums (n > 200). **(e)** Analysis by Uniform Manifold Approximation and Projection (UMAP) showing well-separated clustering of the 15 barcoded spectral vectors from all pooled nanobeads. **(f)** Confusion matrix showing the classification accuracy for molecular multiplexing and decoding.

### Spectrum-CODE based molecular sensing

The Spectrum-CODE system for molecular sensing includes high-throughput individually labelled beads with unique fluorescence code, which were further decoded by machine learning algorisms for quantification. By functionalizing encoded fluorescent beads with antisense oligoes, complimentary to the RNA targets, the beads will specifically hybridize with the RNA bound to the nanoprobes. In this way, we can achieve high throughput detection, identification, and quantification of the RNA targets (**Supplementary Note 2**) by counting the number of the encoded fluorescent beads.

After ‘molecular fishing’, despite thousands of targets on the nanoprobe surface, the beads may not bind all the targets due to the stereo-hindrance effect. The binding events happen in a stochastic manner, where higher targets intensity on the vertical probe array surface will induce higher binding probability (**Supplementary Figure S11, Supplementary Note 3**). To characterize the detection limit of Spectrum-CODE based molecular sensing technique, we first performed a mock experiment by profiling miRNAs (synthetic let-7a) or m^6^A-mRNAs (synthetic m^6^A-RNAs, 31 bp, single-stranded, one m^6^A site) from medium containing a premixed targets at different concentrations varying from 10^-16^ to 10^-10^ M using functional silicon vertical probes array and Spectrum-CODE beads. The silicon vertical probes were functionalized with p19 or m^6^A antibody for miRNAs or m^6^A-mRNAs profiling, respectively. After fishing targets from the medium, Spectrum-CODE with specific reporter probe of the mock targets were utilized for quantification (**Figure 3a**).

**Figure 3.**
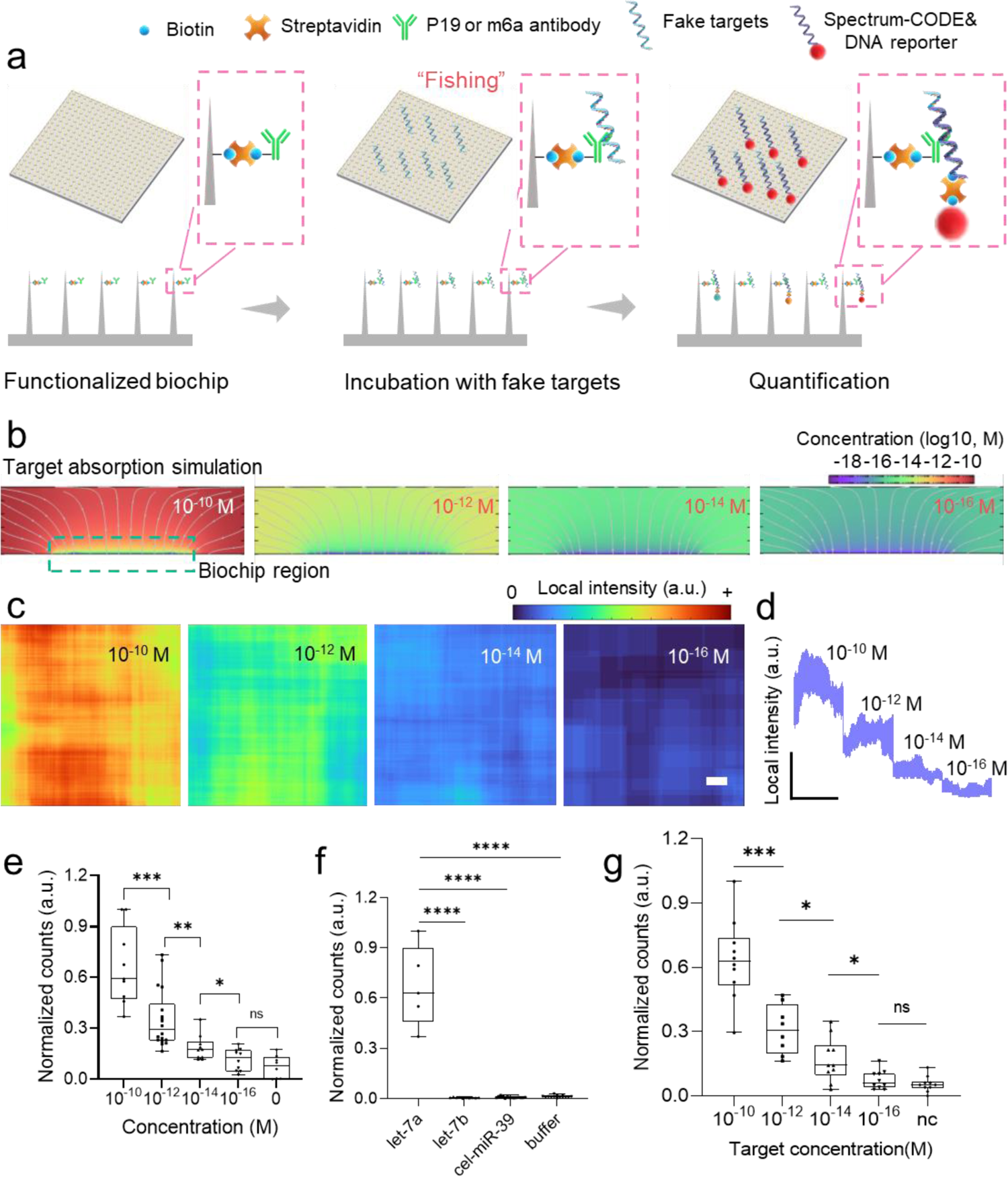
Spectrum-CODE based molecular sensing. **(a)** Schematic of Spectrum-CODE for quantification of molecules extracted by ‘molecular fishing’. **(b)** Molecular targets transport and adsorption simulation performed with COMSOL in different concentration condition. The kinetic equation is based on the Langmuir approximation for the adsorption and desorption rates. **(c)** Representative Spectrum-CODEs beads intensity heatmap on biochip in various concentration. Scale bar, 20 μm. The local intensity was calculated based on the raw bead distribution image by replacing the value of each pixel with the beads counting number in the surrounding 200 pixels square window region. **(d)** To further visualize the beads intensity variation, we transform the 2-dimensional intensity map in **(c)** to 1-dimensional beads intensity curve. **(e, f)** Characterization of the miRNA detection sensitivity and specificity. (Unpaired t test, N>8, (left), N>5 (right); **** p<0.0001; *** p<0.001; ** p<0.01; * p<0.05). **(g)** Characterization of the detection sensitivity for m^6^A-mRNAs by using Spectrum-FISH. (Unpaired t-test, N>8, *** p<0.001; * p<0.05)

For starter, we perform a transport and adsorption simulation using COMSOL. The kinetic equation is based on the Langmuir approximation for the adsorption and desorption rates. With the simulation, we found that the target surface intensity absorbed on the biochip is approximately linear with the solution target concentration, which is also in accordance with our previous finding ^22, 24^ (**Figure 3b**). With the target gradient surface intensity, we can conclude that higher target surface intensity will present higher binding probability of the Spectrum-CODE beads. To validate this assumption, we perform the mock experiment with gradient concentration. Representative visualization of the beads binding intensity with different concentration shows a recognizable difference and relatively homogeneous spatial distribution (**Figure 3c, d, Supplementary Figure S12**). Considering systematic variations, more replications were further demonstrated and the binding numbers in different batches of mock experiments were summarized (**Figure 3e, f, g**). The 10^-10^ M vs 10^-12^ M, 10^-12^ M vs 10^-14^ M and 10^-14^ M vs 10^-16^ M all presents significant difference in both miRNA and m^6^A mRNA mock experiment, particularly our platform presents high specificity with even 2-nt mismatches (let 7a vs let 7b) in the miRNA mock experiment (**Figure 3f**). The above test further indicate that the detection limit can reach as low as 10^−14^ to 10^-16^ M^23^ (corresponding to ∼6000 copies/μL to ∼60 copies/μL), which is much lower than the abundance of even a single copy in a cell (considering the cell as a 20 μm diameter sphere, the concentration for single copy is ∼10^-13^ M).

### Tissue-wide single cell profiling of miRNAs

The Spectrum-FISH technique was then applied to examine acute OB tissue slices to demonstrate a tissue-wide single cell spatial profiling of miRNAs. To correlate the resulted post-transcriptional miRNA profiles with the spatial distribution of individual cells, a piece of acute OB tissue was sectioned into two opposing slices, one was immunostained for cell-type specific markers (i.e., GFAP for astrocytes and NeuN for neurons) and the other was examined by Spectrum-FISH miRNA analysis. The results from the two opposing slices were then registered to give a full picture of the heterogeneous post-transcriptional miRNA regulation in the coordinates of all identified cells (**Figure 4a**). The expression of 24 targeted miRNAs (**Supplementary Table S1**) were analyzed. To accommodate such a throughput, three rounds of microscopic visualization were performed (**Supplementary Figure S5**). In each round, a double stranded non-mammalian miRNA (cel-miR-39) was introduced to serve as an internal reference for data normalization, thus eliminating interference from systematic variations. For individual nanoprobes, the expression of extracted miRNAs was quantified by counting the beads of specific spectrum codes (**Figure 2c**). For a particular type of cells, such as the GFAP^+^ astrocytes, the vertical nanoprobes that were in contact with each cell were respectively identified (**Figure 4b-d**). For an individual cell, the subcellular distribution of particular miRNAs can be directly visualized by the distribution the nanobeads of corresponding spectrum code (**Figure 4e-g**). From all nanoprobes under the umbrella of a specific cell, the number of miRNA-specific nanobeads were summed up and used as a quantification of the cell’s overall expression. When the single-cell miRNA data was pooled together for an informatic analysis, different astrocyte groups (ast1-ast5) were revealed by unsupervised consensus clustering (**Figure 4h-j, Supplementary Figure S13**). While the biological basis for the identified astrocytes subpopulations is not clear, our results directly showed the spatially distributed heterogeneity of post-transcriptional regulation in a particular cell type. Similar observation of neuronal subpopulations could also be derived by miRNA heterogeneity analysis in NeuN^+^ cells (**Supplementary Figure S14**).

**Figure 4.**
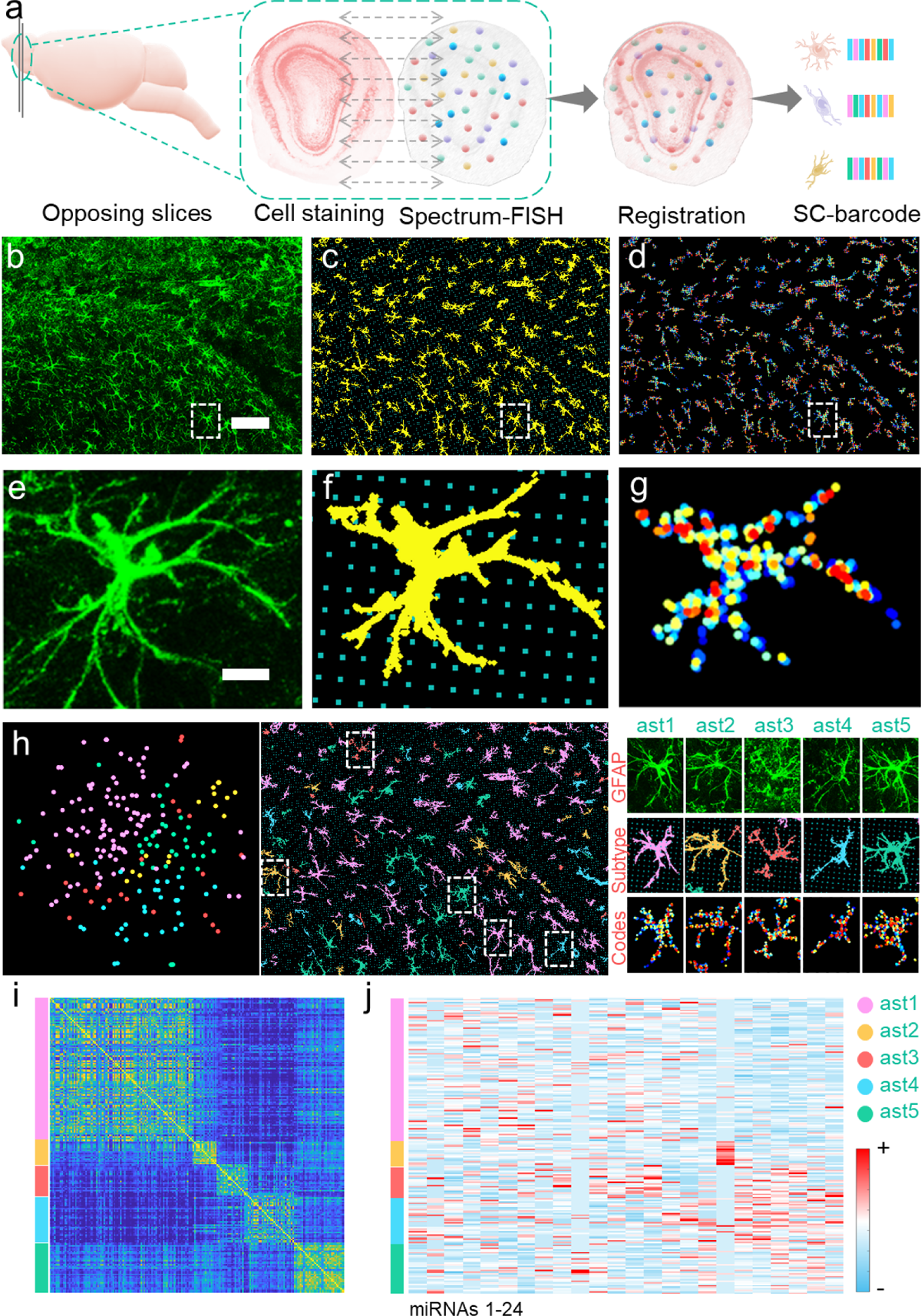
Tissue-wide single cell profiling of miRNAs. **(a)** Schematic of the experimental procedures that correlate Spectrum-FISH miRNA analysis with cell-type specific (GFAP for astrocytes and NeuN for neurons) immunostaining. The results from two opposing slices were registered to give a full picture of the heterogenous post-transcriptional miRNA regulation in the coordinates of all identified cells. **(b)** Fluorescence image for GFAP staining of an OB slice. Scalebar, 100 μm. **(c)** Spatial registration of identified astrocytes (GFAP^+^) with the nanoprobes that has been used for Spectrum-FISH. **(d)** Quantification of nanoprobe-associated nanobeads with miRNA-specific spectrum codes (in different colors). **(e-g)** Enlarged view of the boxed region in panel **b-d**. Scalebar, 10 μm. **(h)** Analysis of cellular subtypes in GFAP^+^ cells using unsupervised consensus clustering of the miRNA vectors. The left column shows the t-SNE distribution of the identified subtypes; the middle column shows the spatial distribution of different astrocyte subtypes in association with the nanoprobes; the right column shows representative the cellular morphology and the miRNA-specific spectrum code distribution for each astrocyte subtype. **(i)** Consensus matrix generated from the consensus clustering analysis using pooled single-cell miRNA data. **(j)** The expression heatmap showing the expression of 24 miRNAs in the GFAP^+^ cells and in the identified astrocyte subtypes.

### Spatial heterogeneity of miRNA regulation in olfactory bulb

To fully unleash the power of Spectrum-FISH for tissue-wide post-transcriptional profiling, we then analyzed the spatial expression of 24 miRNAs across a whole coronal OB slice. An OB slice were labelled by six anatomical regions based on the Allen Brain Atlas (**Figure 5a**)^25^, to which the spatial correlation of the nanoprobe-derived miRNA vectors were explored. For an OB slice as large as ^∼^3 mm^2^, signal from ∼110000 nanoprobes was examined. In our computational analysis, the nanoprobes was processed with 4×4 binning (**Supplementary Figure S15**), which effectively divided an OB slice into ^∼^6800 regions of interest (ROIs, ^∼^24×24 µm^2^/each). From each ROI, a 24-dimension miRNA vector was derived by the expression of the targeted 24 miRNAs. The miRNA vectors from different OB regions were first examined by uniform manifold approximation and projection (UMAP), which clearly showed relatively more coherence for vectors in close spatial proximity (**Figure 5b**). Our analysis also showed statistical similarity in miRNA expression among different OB regions, such as outer plexiform layer (OPL) *vs.* mitral layer (ML), granule layer (GR) *vs.* inner plexiform layer (IPL) or outer plexiform layer (OPL). The miRNA expression in the glomerular layer (GL) or subependymal zone (SEZ) showed more distinctive patterns (**Figure 5c**). The observed miRNA coherence could be reasonably explained by the spatial distance of neighboring OB regions, it also echoes previous reports that documented similar cellular composition in OPL, ML, IPL and GR ^26^.

**Figure 5.**
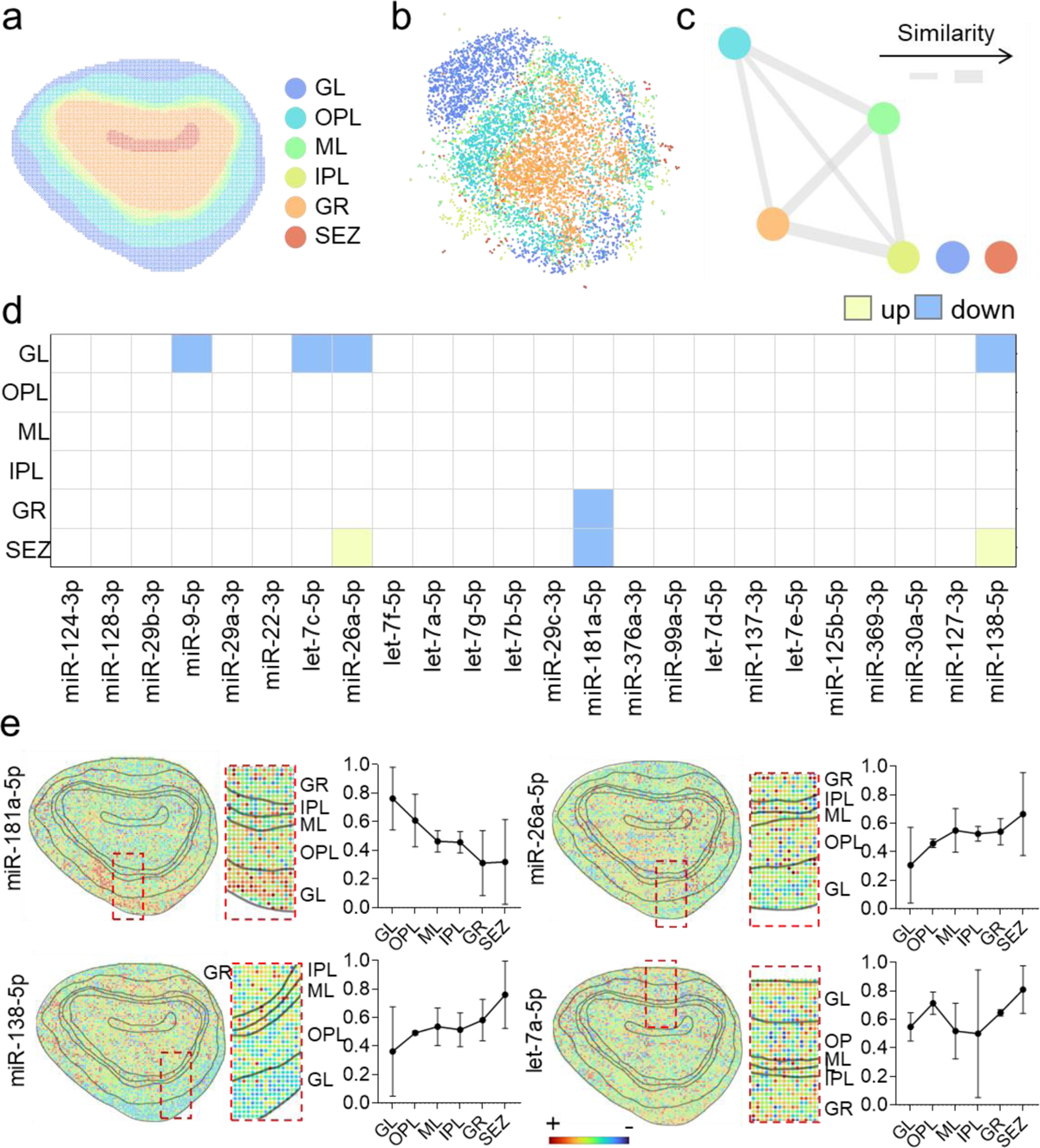
Spatial distribution of miRNA expression in different olfactory bulb (OB) regions. **(a)** Labeling of six anatomical regions in a coronal OB slice. **(b)** Coherence analysis of the miRNA vectors from different OB regions by using uniform manifold approximation and projection (UMAP). **(c)** The similarity network of the six OB regions derived by using the associated miRNA vectors. Shorter distance and thicker connection represent higher similarity in miRNA expression. **(d)** Identification of the dominant miRNA signature (out of the 24 targets) for each OB region. The comparison was made between one region versus all others, a signature was defined as a fold change greater than 1.1 and a p<0.05 by unpaired t-test. **(e)** Heatmap showing the representative spatial expression of a particular miRNA across a whole OB slice. The analysis was performed with a 4×4 binning of the nanoprobes, the error bars indicate mean ± SD, data from more than 300000 nanoprobes were collected from three biological replicates.

To identify the dominant miRNA signature (out of the 24 targets) for each OB region, the expression of a particular miRNA (average from all associated nanoprobes) was statistically compared across the six OB regions (**Figure 5d**). It was found that miR-181a-5p was down-regulated in GR and SEZ, and there is a general decreasing expression level of miR-181a-5p across the outer to inner region of OB. Oppositely, miR-138-5p showed significant up-regulation for the most inner part of OB in the SEZ layer, and an increasing expression pattern was observed across GL to SEZ regions; similar trend was also found with miR-26a-5p (**Figure 5d, e**). Interestingly, let-7a-5p show a relatively even distribution across different OB regions, suggesting different roles of particular miRNAs for a heterogeneous spatial regulation in the OB ^27^.

To further explore the intrinsic spatial miRNA patterning, unsupervised BayesSpace ^28^ clustering analysis was performed without prior OB structural labelling. As shown in **Figure 6a, b**, all the miRNA vectors acquired from the ^∼^6800 ROIs were identified to group into 8 clusters solely based on the miRNA expression coherence. The spatial distribution of these clusters was identified to show potential enrichment tendency in particular OB anatomic regions (**Figure 6c**), which was statistically confirmed by a hypergeometric test (**Figure 6d**). For example, miRNA cluster 2 (miR-C2) and miR-C5 were enriched in GL (**Supplementary Note 4**). For each identified miRNA clusters, the most phenotypic signature miRNAs were then determined (**Figure 6e**). From these analyses, it is likely that some miRNAs are particularly important for the spatially heterogeneous post-transcriptional regulation in OB, especially in the region of OPL and GR (**Figure 6f, Supplementary Figure S16**).

**Figure 6.**
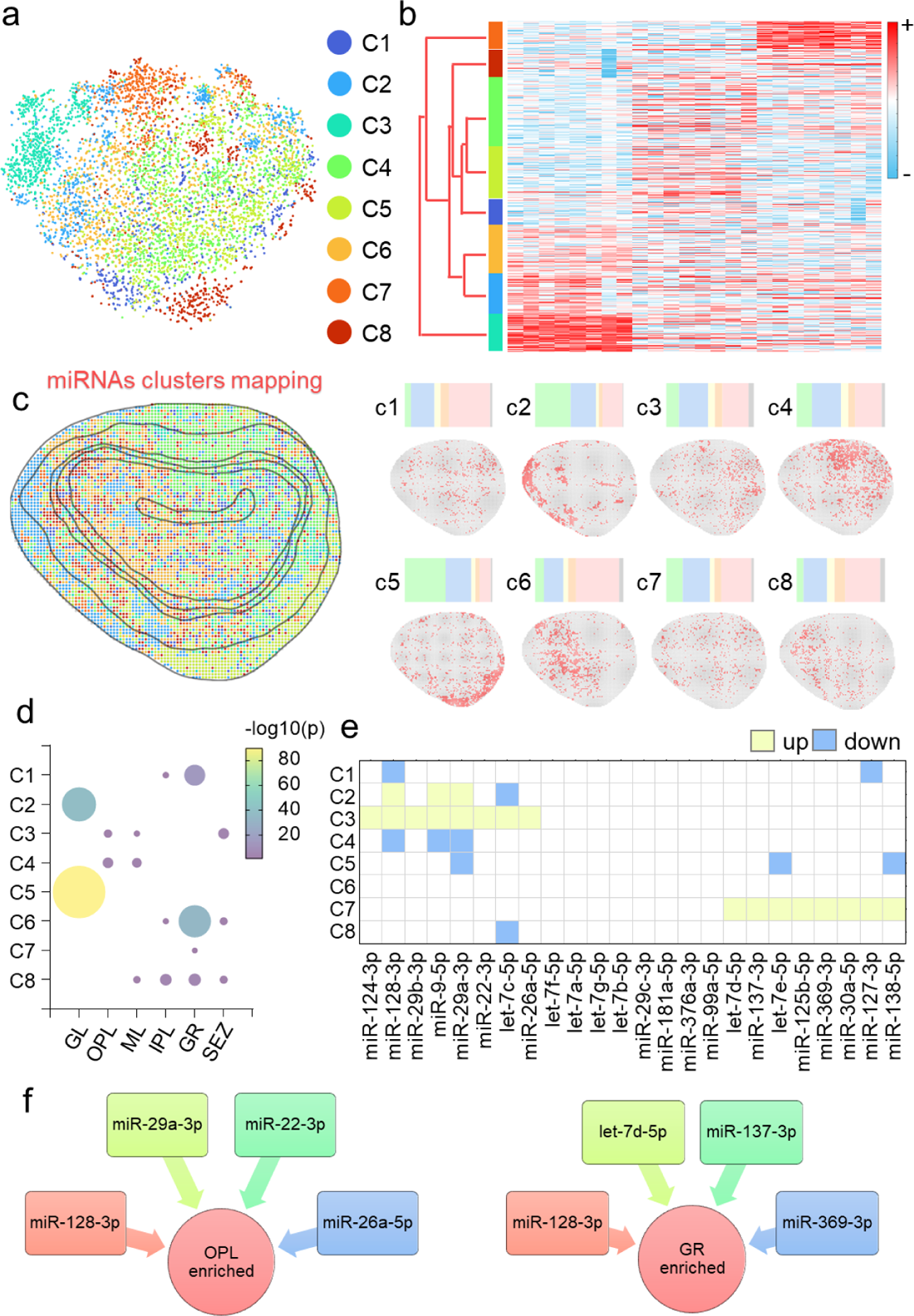
Unsupervised clustering analysis of miRNA spatial heterogeneity in OB. **(a)** t-SNE visualization of the eight miRNA clusters derived by unsupervised BayesSpace clustering. **(b)** Heatmap showing normalized miRNAs expression of the 8 clusters with the dendrogram tree indicating their hierarchical similarity. **(c)** Mapping of the spatial distribution of 8 miRNA clusters in an OB structure atlas. The split subpanels (right) show the mapping of individual miRNA clusters and the corresponding enrichment in different OB sub-regions. **(d)** Visualization of the enrichment analysis of the 8 clusters in six OB sub-regions by using hypergeometric test. The dot size and color indicate regional enrichment, which is derived from the P value of the hypergeometric test. **(e)** Identification of the miRNA signature (out of the 24 targets) for different miRNA clusters. The comparison was made between one cluster versus all others, a signature was defined as a fold change greater than 1.5 and a p<0.05 by unpaired t-test. **(f)** Visualization of the potential correlation between the miRNA cluster signature and spatial enrichment in specific OB sub-regions (OPL and GR). Data from more than ^∼^300000 nanoprobes were collected from three biological replicates.

### Spatial regulation of RNA methylation in olfactory bulb

In addition to miRNAs, the Spectrum-FISH can be applied to analyze other post-transcriptional targets (e.g. RNA methylations), so that the spatial cooperative involvement of different post-transcriptional mechanisms can examined. Towards this goal, we then used Spectrum-FISH to investigate the spatial dynamics of 9 mRNAs with m^6^A methylation in an acute coronal OB slice (**Supplementary Table S2**). The “molecular fishing” system was adapted by replacing the “bait” protein (p19 for miRNAs) with m^6^A-specific antibodies, and the nanoprobe associated experimental operations remained the same. The m^6^A-specific antibodies can extract all m^6^A-methylated RNAs, which were later decoded by an on-chip analysis and quantification (**Supplementary Figure S17)**.

The expression of the 9 m^6^A-mRNAs in different OB sub-regions was firstly examined, showing a unique pattern in SEZ in comparison to other OB regions (OPL, ML, IPL and GR) with significant upregulation of m^6^A-Gpr161 and down-regulation of m^6^A-Epha7 in SEZ. Such observation was further confirmed by a spatial mapping of these m^6^A-mRNAs (**Supplementary Figure S18**). Using unsupervised clustering analysis, the 9-dimensional m^6^A-mRNA vectors (from all nanoprobes covered by an OB slice) was self-grouped into 8 clusters (m^6^A-C1 ∼ m^6^A-C8, **Figure 7a-c**), of which m^6^A-C4 and m^6^A-C7 were particularly enriched in SEZ. Meanwhile, with a moderate level, m^6^A-C2 was enriched in OPL, m^6^A-C5 & m^6^A-C7 were enriched in GL, and m^6^A-C6 was enriched in GR (**Figure 7c, d**). These results further show a substantial heterogeneity in mRNA methylation at different OB regions, suggesting a complex role of associated translational regulation of cellular function (**Supplementary Figure S19**).

**Figure 7.**
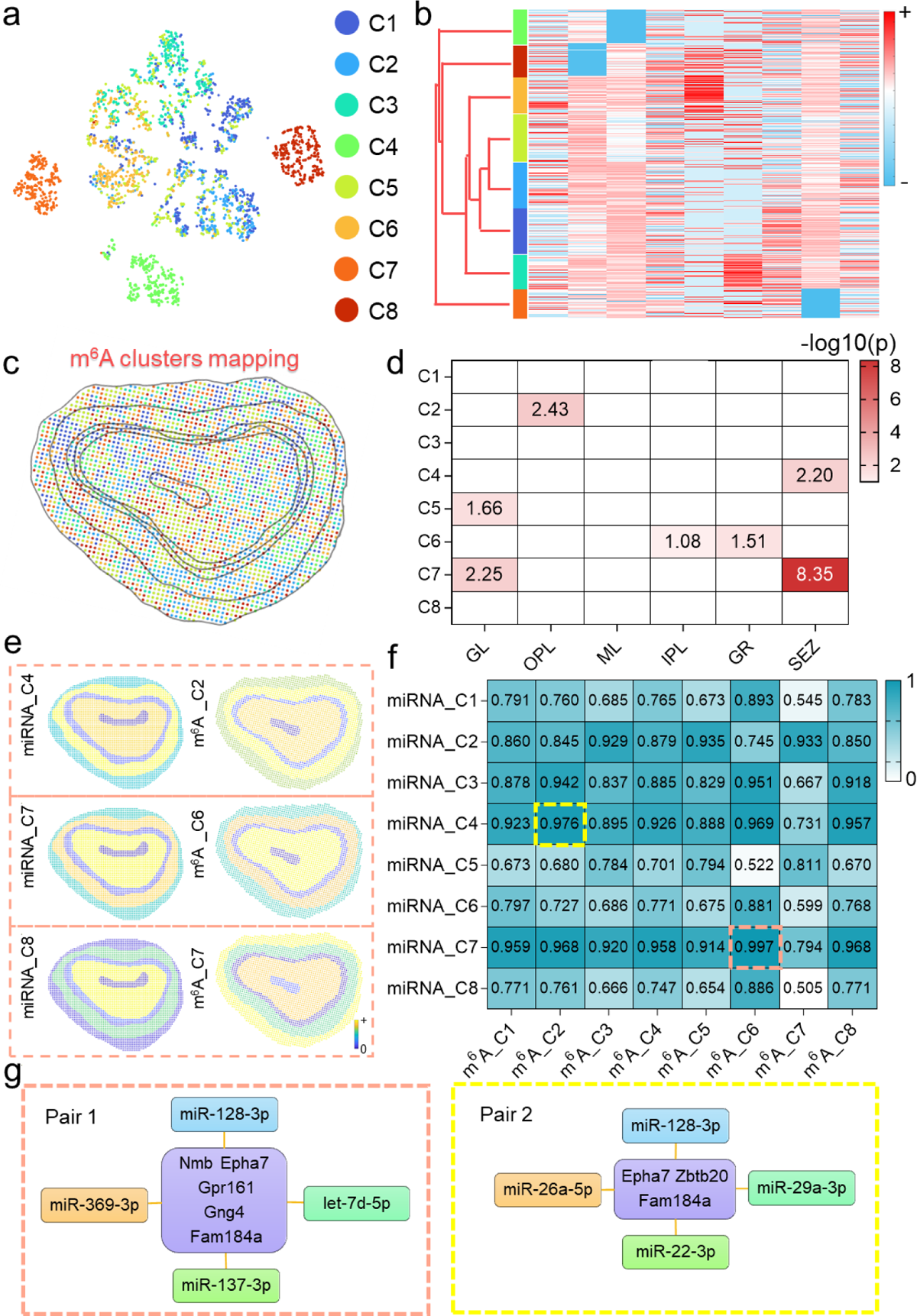
Tissue-wide spatial heterogeneity of RNA methylation in OB. **(a)** t-SNE visualization of the eight m^6^A-mRNA clusters derived by unsupervised BayesSpace clustering. **(b)** Heatmap showing normalized expression the 9 m^6^A-mRNAs for the 8 identified m^6^A-mRNA clusters with the dendrogram tree indicating their hierarchical similarity. **(c)** Mapping of the spatial distribution of 8 m^6^A-mRNA clusters in an OB structure atlas. **(d)** Analysis of the spatial enrichment of each m^6^A-mRNA clusters in six OB sub-regions by using the hypergeometric test. **(e)** Visualization of representative spatial distribution vectors (SDV) for coupled miRNA and m^6^A-mRNA clusters. **(f)** Analysis of spatial correlations between paired miRNA- and m^6^A-mRNA clusters with Pearson correlation coefficient **(g)** Identification of potential cooperative involvement of paired miRNAs and methylated mRNAs in different OB sub-regions.

As previously reported, m^6^A methylation of mRNAs can be regulated by miRNAs via a sequence pairing mechanism to modulate the binding between methyltransferase and mRNAs ^18^, the versatility of the Spectrum-FISH technique provides an extra dimension to study the cooperative involvement of the two post-transcriptional regulatory mechanisms across a whole tissue sample. A spatial distribution vector (SDV) for miRNA clusters or m^6^A-mRNA clusters were generated (**Figure 7e**), which were used to derive the spatial correlations between paired miRNA- and m^6^A-mRNA clusters (**Figure 7f**). Notably, miRNA-C7 and m^6^A-C6 pair showed the highest spatial correlation, and both clusters were identified to enrich in GR (**Figure 6d, 7d**), indicating a high level of cooperatively of associated miRNAs and m^6^A methylation in the region. Another highly correlated pair was miRNA_C4 and m^6^A_C2, which both showed enrichment in OPL (**Figure 6d, Figure 7d**). Considering the signature targets in different miRNA- or m^6^A-clusters, it would be interesting to further explore the potential interactive roles of paired miRNAs and methylated mRNAs (**Figure 7g**), especially in GR or OPL region of rodent OB.

## Discussion

While the importance of genetic or epi-genetic heterogeneity has been increasingly recognized^21, 29, 30^, it has been challenging to profile intracellular post-transcriptional targets with sufficient throughput and resolution, especially when such analysis is needed at a large-scale across centimeter wide tissue samples ^15, 21^. Current developed spatial profiling techniques, either FISH or sequencing based, mostly can only be used to profile mRNAs and very limited techniques ^8, 16, 17^ can profile other type of targets (Proteins&mRNAs^8, 16^, chromatin modifications^17^). Towards this goal, we describe a technique, Spectrum-FISH, which enables high-throughput tissue-wide spatial profiling of various post-transcriptional targets in acute tissue sections with single cell resolution. The platform uses a biochip with an array of vertically aligned nanoprobes to effectively extract intracellular molecules (miRNAs, m^6^A-mRNAs) for downstream analysis in the coordinates of the large-scale of cells within a tissue slice and shows potential correlation among replicative bio-samples (**Supplementary Note 4, Supplementary Figure S20-S22**). In our previous works, diamond nanoprobe-assisted molecular fishing have been previously employed for analyzing miRNAs in living cells or fresh tissue samples ^22^, this study, for the first time, achieved a spatial post-transcriptome analysis of both miRNAs and methylated mRNAs with single cell resolution across a millimeter to centimeter brain tissue slice.

Besides the expression information, the key for a spatial molecular profiling is a feasible strategy to preserve the microscale coordinate information in association with the molecular expression levels, which is currently the bottleneck for most efforts in developing spatial transcriptome techniques ^8, 9, 16^. In our Spectrum-FISH, the molecular information is sampled and preserved by individual nanoprobes, which are registered to thousands of cells in an acute tissue slice after the initial cellular contact for intracellular molecular fishing. To achieve such spatial mapping, a UV imprinting scheme was designed to acquire a pixelated tissue morphology feature on the array of nanoprobes with larger footprint than the tissue. As all the nanoprobes were labeled with a photocleavable fluorescent dye, a brief UV exposure was sufficient to cleave off the dyes on nanoprobes that were not covered by the tissue sample, thus to generate a contrasted fluorescent pattern of the tissue outline on the chip, providing a reference framework for spatial registration to later tissue images acquired by immunostaining and optical microscopy. Unlike existing spatial indexing methods that typically involves molecular sampling with tremendous efforts in spatial control ^8, 9, 16^, our UV imprinting facilitated spatial encoding is easy to implement and does not require any special equipment, such as robotic sampler or microfluidic chamber, making it extremely cost-effective. Together with a custom-made imaging processing algorithm, each nanoprobes can be traced back to individual cells at subcellular resolution to reveal the post-transcriptional profiles across a whole centimeter tissue sample.

Another challenge for imaging based spatial transcriptome analysis is the limited numbers of fluorescent channels that restricting the multiplexing throughput. Previously, FISH-based method and NanoString barcoding have been reported. FISH based fluorescence encoding in some cases^5–7^ requires super-resolution microscopy and involves serial rounds of hybridization, imaging and probe stripping ^5, 6^, making it expensive, time-consuming and technically difficult for wider adoption. For NanoString barcoding, the barcodes were prepared using a string of RNA particles coupled with fluorophores, which requires intricate sequence design and synthesis processes. In an assay, the reporter barcodes also need to be stretched by an electric field before imaging, further limiting its usage for in situ analysis. Though NanoString has been used in a spatial mRNA profiling study ^16^, the implementation was of relatively low spatial resolution and slow profiling speed due to the sequential region by region cleaving process of the reporters.

Alternatively, for the Spectrum-FISH technique, we developed a spectrum digitization barcoding method for functional differentiation of multiple analytical targets, which relies on a rainbow fluorescence composition and machine learning-based spectrum decoding. The simplest spectrum digitization method uses a “1” (with dye) or “0” (without dye) mixing to compose the spectral codes. Specifically, for analyzing RNAs, a DNase-assisted multi-round visualization process ^6^ further expand the code pool. As demonstrated here, we used a four-channel system to encode 27 miRNAs, and the machine learning based spectrum decoding gives more than 90% accuracy. Theoretically, the encoding pool can be further expanded if the rainbow mixing changes from the current binary format and adopts a step-wise ratio for different fluorophors. Accordingly, the number of available codes will be *N* × (R_step_^C^ − 1), where R_step_ indicates the ratio step number, *C* indicates the fluorophore channels number, and *N* indicates the rounds of visualization cycles. In a trinary system with 7 different fluorophores would substantially increase the multiplexing throughput to over 10,000 by 5 rounds of visualization (**Supplementary Methods**).

In most existing spatial transcriptomic technologies, tissue samples require extensive pre-treatment to release mRNAs, which could affect the analytical resolution by cross contamination among neighboring regions. Spectrum-FISH, on the other hand, directly applies to freshly prepared acute tissue slices without any further processing, thus ensuring almost instant detections. The nanoprobe-based in situ analysis also help avoid the decomposing and reassembly processes that are commonly required in existing methods. Moreover, existing spatial profiling techniques are mostly applicable to a single type of molecules, mRNAs or proteins ^8, 16^, Spectrum-FISH is also advantageous in its applicability towards different targets by a customized functionalization of molecular “baits” on the nanoprobes, such as p19 protein for miRNAs or antibody for m^6^A-mRNAs, as demonstrated in this study. Such a versatility can be readily extended to other ‘baits’, such as poly(T) sequences for mRNAs, different antibodies for signaling proteins or methylated RNAs, even RNA-binding proteins (RBPs) for interactive translational regulation factors. The multi-omics spatial heterogeneity and correlation of different post-transcriptional regulations remains largely unexplored, which could be potentially resolved by our Spectrum-FISH technique, as illustrated by our discovery of highly associated miRNAs and m^6^A-mRNAs in specific OB regions. While the exact mechanism of the interactive regulation between the identified miRNAs and m^6^A-mRNAs remain to be elaborated, these results show a proof-of-concept investigation of the spatial dynamics of multiple post-transcriptional targets in tissues of millimeter to centimeter scales. It is worth to notice that the miRNAs or m^6^A-mRNA profiling was not conducted simultaneously in this study, such multi-omics spatial epi-transcriptome analysis can also be explored by a “dual-bait” functionalization of the nanoprobes, so that the interactive roles of miRNAs and specific methylated mRNAs can be further explored with more delicate specificity and spatial dynamics, which could potentially provide new insights to the progress and treatment of human diseases ^29–31^.

## Methods

### Nanoprobe functionalization

Silicon nanoprobe chips were prepared using deep Reactive Ion Etching (DRIE), followed by thermal wet oxidization and chemical etching steps^32^. Silicon nanoprobe chips were cut into squares (5 mm×5 mm) and then processed with piranha solution (v/v = 3:1, 98% H_2_SO_4_:27.5% H_2_O_2_) at 90 ℃ for 90 min. The chips were subsequently washed with Deionized water, methanol, methanol/dichloride methane (DCM) mixture (v/v = 3:1) and DCM. (3-aminopropyl)triethoxysilane with a volume fraction of 3% in DCM was used to further functionalize the chips with amino functional groups for 3 hours. Afterward, the chips were washed by ethanol, isopropyl alcohol and deionized water subsequently.

The amino functionalized chips were activated using glutaraldehyde (15%, v/v) for 2 hours and then crosslinked with p19 siRNA binding protein (1 μg/ml in depc-PBS; New England Biolabs) or anti-N6-methyladenosine (m^6^A) antibody (1 μg/ml in depc-PBS, Abcam) for 2 hours. BSA (1%, m/m in depc-PBS) and triton X100 (0.1%, v/v, in depc-PBS) mixture were used to block the unreacted groups of the chips for 5 hours. The chips were further reacted with PC biotin-PEG3-NHS ester (0.2 mg/ml, Sigma-Aldrich) for 1 hour and then then labeled with streptavidin conjugated with Alexa Fluor^TM^ 568 (0.04 mg/ml in depc-PBS, ThermoFisher) for 1 hour, followed with treating with unlabeled biotin (0.1 mg/ml) for another 2 hours to block the unreacted streptavidin sites. All the reactions were performed in room temperature unless specifically mentioned.

### Spectrum codes fabrication

50 μL amino magnetic beads (J&K, 300-400 nm diameter, in PBS) were used for the spectrum codes preparation followed by reacting with 1 μL biotin-NHS (Sigma, 2 mg/mL) for 2 hours in room temperature. Then the beads were washed 3 times using depc-PBS and 5 μL diverse type of streptavidin conjugated with specific fluorophores (i.e. Streptavidin, Alexa Fluor™ 488 conjugate, Streptavidin, Alexa Fluor™ 514 conjugate, Streptavidin, Alexa Fluor™ 555 conjugate, Streptavidin, Alexa Fluor™ 647 conjugate, ThermoFisher, 2 mg/mL) were mixed together with biotin functionalized beads for 2 hours in room temperature. Depc-PBS was used to wash the beads 3 times and 4 μL corresponding biotin DNA probe (BGI, 100 μM) was mixed with the beads for 2 hours, followed with treating with unlabeled biotin (0.1 mg/ml) for another 2 hours to block the unreacted streptavidin. After washing three times with depc-PBS, the beads were dispersed in 50 μL 1% BSA and 0.1% trition X100 solution and stored in 4 ℃ for further applications.

### Spatial probing of miRNAs via intracellular biopsy

All the complementary miRNA probes with overhang were mixed with a concentration of 10^-8^M in hybridization buffer (1%, m/m, BSA, 0.01%, v/v, tween 20, 5×SSC), followed by adding miR-cel 39 (10^-10^M) as internal reference and hybridize with the complementary probe for 15 min in 37 ℃ water bath.

The experiments involving animals are approved by the Ethics Committee of City University of Hong Kong. C57BL/6 mice (4-6 weeks) were sacrificed by cervical dislocation. The brains were harvested and maintained in ice-cold depc-PBS buffer, mounted on a vibratome with glue, and then were cut into coronal slices each of 500 μm thick.

Acute brain slice was rinsed with depc-PBS for several times and then transfer under the needle chip immersed with the prepared probes in a four well plate. The plate was centrifuged at 500 rpm (35.5 g of RCF, same below) for 5 min to initiate a membrane puncture. The slices (or cells) with the nanoprobe patch were incubated 15min before miRNA target extraction in 37 ℃. The sample again underwent a centrifugation at 500 rpm for 10 min to fish targeted miRNAs from brain slices (or cells) for further analysis.

After intracellular fishing, the needle chip with tissue on the top was transferred in depc-PBS with 0.1% tween 20 and exposed to 5 mW/cm^2^ UV for 20 min to imprint the tissue outline. A digital photograph was also captured to record the tissue and needle spatial relative location as an auxiliary registration method complementary to the UV imprint. Then the tissue was removed from the needle chip. Barcodes with different reporter probes were used to label the targets in the needle chip for 2 hours. Leica SP8 with 63×oil objective was used for imaging.

### Spatial probing of m^6^A mRNAs via intracellular biopsy

Acute brain slice was rinsed with depc-PBS for several times and then transfer under the needle chip immersed with the prepared probes in a four well plate. The plate was centrifuged at 500 rpm (35.5 g of RCF, same below) for 15 min to fish targeted m^6^A mRNAs from brain slices (or cells) for further analysis.

After intracellular fishing, the needle chip with tissue on the top was transferred in depc-PBS with 0.1% tween 20 and exposed to 5 mW/cm^2^ UV for 20 min to imprint the tissue outline. A digital photograph was also captured to record the tissue and needle spatial relative location as an auxiliary registration method complementary to the UV imprint. Then the tissue was removed from the needle chip. Barcodes with different reporter probes were used to label the targets in the needle chip for 2 hours. Leica SP8 with 63×oil objective was used for imaging.

### Strip and re-hybridization

After finished imaging of one round, DNase I (ThermoFisher) was used to digest the reporter probes and strip the barcodes in 37 ℃ for 30 min. The needle chip sonicated in triton X100 (0.25%, v/v, in depc-DI) for 30 s and washed with depc-PBS for 3 times. Another round barcode with reporter probes were used to label the targets of the needle chip.

### Image processing and analysis

Confocal microscope (Leica SP8) equipped with 63× oil-immersion objective and Leica Application Suite X (LAS X) software was used for imaging and data collection. Registration of tissue and needle array were performed in Photoshop and MATLAB R2021a image processing toolbox using the UV imprint outline, digital photo and the tissue immunofluorescence staining image. The labelled brain region of the staining image was performed in Photoshop based on Allen brain atlas. For the single cell analysis, watershed segmentation in ImageJ-1.53c was used to process the NEUN stained image. The segmentation of the GFAP stained image was performed by adaptive binarization in MATLAB R2021a. For decoding the barcodes, a threshold was optimized for the fluorescence image binarization. ROIs were extracted to generate barcode feature vectors for the further decoding using ML model. The ML was performed using classification toolbox in MATLAB R2021a, where ensembled bagged trees model was used with a learner type of decision tree. The maximum number of splits is 22087 and 30 learners were applied.

### Statistics analysis

For Fig. 1, an olfactory bulb brain slice tissue with ∼3 mm^2^, signals from ∼110000 nanoprobes were examined and presented as a technique demonstration. For quantitative analysis, at least three biological replicates were used in this study. For Fig. 2, the fluorescence vectors and associated barcodes were acquired from 15 types of nanobeads with different digital spectrums (More than 200 replicates were randomly selected for each type of nanobeads). For Fig. 3, more than 8 replications were used for characterization of the miRNA detection sensitivity and more than 5 replications were used for the miRNA detection specificity. In characterization of the detection sensitivity for m^6^A-mRNAs, more than 8 replications were analyzed. For Fig. 4, a rectangular region with ∼0.45 mm^2^ was randomly selected for single cell profiling demonstration. ∼198 astrocytes (GFAP^+^) were identified and used for consensus clustering. The elbow method was used to determine the optimal K based on the CDF curve. 5 subtypes with distinct expression pattern were discovered. For Fig. 5, to identify the dominant miRNA signature (out of the 24 targets) for each olfactory region, the comparison was made between one region versus all others and a signature was defined as a fold change greater than 1.1 and a p <0.05 by unpaired t-test. The analysis was performed a 4×4 binning of the nanoprobes, the error bars indicate mean ± SD, data from more than 300000 nanoprobes were collected from three biological replicates. For Fig. 6, to identify the miRNA signature (out of the 24 targets) for different miRNA clusters, comparison was made between one cluster versus all others and a signature was defined as a fold change greater than 1.5 and a p<0.05 by unpaired t-test. Data from more than ∼300000 nanoprobes were collected from three biological replicates. For Fig. 7, to identify potential cooperative involvement of paired miRNAs and methylated mRNAs in different OB sub-regions, in total ∼9 groups of miRNA-m^6^A mRNAs pair from 6 different biological tissues were used as replicates.

## Acknowledgements

This work was supported by National Natural Science Foundation of China (81871452, 22078184), by the Science Technology and Innovation Committee of Shenzhen Municipality (JCYJ20170818100342392), by General Research Fund (11278616, 11103921) from the Research Grants Council of Hong Kong SAR, by Health and Medical Research Fund (06172336) from the Food and Health Bureau of Hong Kong SAR, and by Guangdong Basic and Applied Basic Research Foundation (2019B030302012). Support from the Hong Kong Centre for Cerebro-cardiovascular Health Engineering and Funds from City University of Hong Kong (7005084, 7005206) are also acknowledged.

## Author Contributions

P.S. conceived the project and supervised the study. X.J. designed the spectrum codes system, optimized the nanoprobes fabrication process, performed spatial profile experiments and further bioinformatic analysis. P.F., X.Z., R.Y. and Y.W. helped with different aspects of the experiments. C.X. and Q.Y. helped with the fabrication of the nanoprobes. L.H. helped with the experimental design for epigenetic profiling. L.Q and X.W helped with the bioinformatic analysis. Z.W. and W. Z. helped with the validation of the intracellular biosensing principle. X.J. and P.S. wrote the manuscript.

## Data availability

The data that support the findings of this study are available from the corresponding author on request.

## Supporting Information

### Supplementary Figures

**Supplementary Figure S1.**
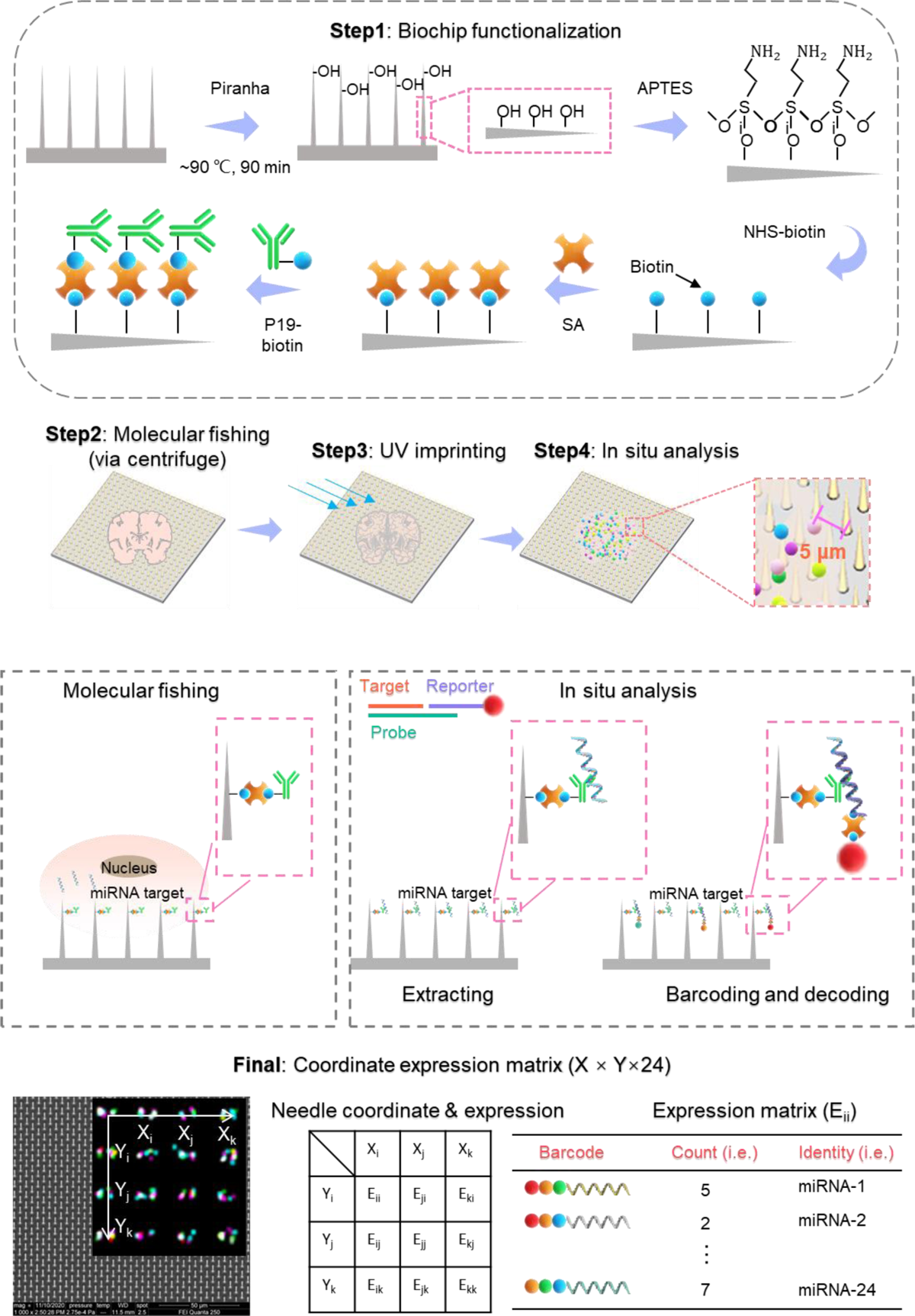
Schematic and procedures of Spectrum-FISH. In the beginning of Spectrum-FISH technique, based on different spatial profiling purpose, the biochip can be functionalized specifically targeting on various types of molecules, such as p19 protein for miRNAs or antibody for m^6^A-mRNAs, as demonstrated in this study. The Spectrum-FISH technique is based on the continuous development of a “molecular fishing” system, which uses an array of silicon nanoprobes as fishing rods for minimum-invasive access of cytoplasmic regions of mammalian cells. Specifically, for “fishing” miRNAs, a size-dependent RNA binding protein, p19, was cross-linked to functionalize the nanoprobes, working as the “fishing hook” to capture double-strand RNAs (dsRNAs). In the Spectrum-FISH system, after the molecular fishing procedure when the tissue was still in contact with the biochip, the spatial encoding was facilitated by a UV imprinting process, which gives a snapshot of the tissue morphology and boundary on the nanoprobe biochip and provides an associative reference linking the cell and nanoprobe spatial coordinate. After the UV imprinting procedure, the silicon nanoprobes biochip was separated from the tissue slice with the target molecules extracted by specifically binding with the ‘bait’. For multiplex visualization and quantification of the captured biotargets, we developed a digital spectrum method (Spectrum-CODE) for functional encoding of the analytical targets. A schematic for miRNA capture probe design was also presented, where a probe formed double-strand RNAs with the target and a reporter probe with Spectrum-CODE was complementary to the overhang for quantification.

**Supplementary Figure S2.**
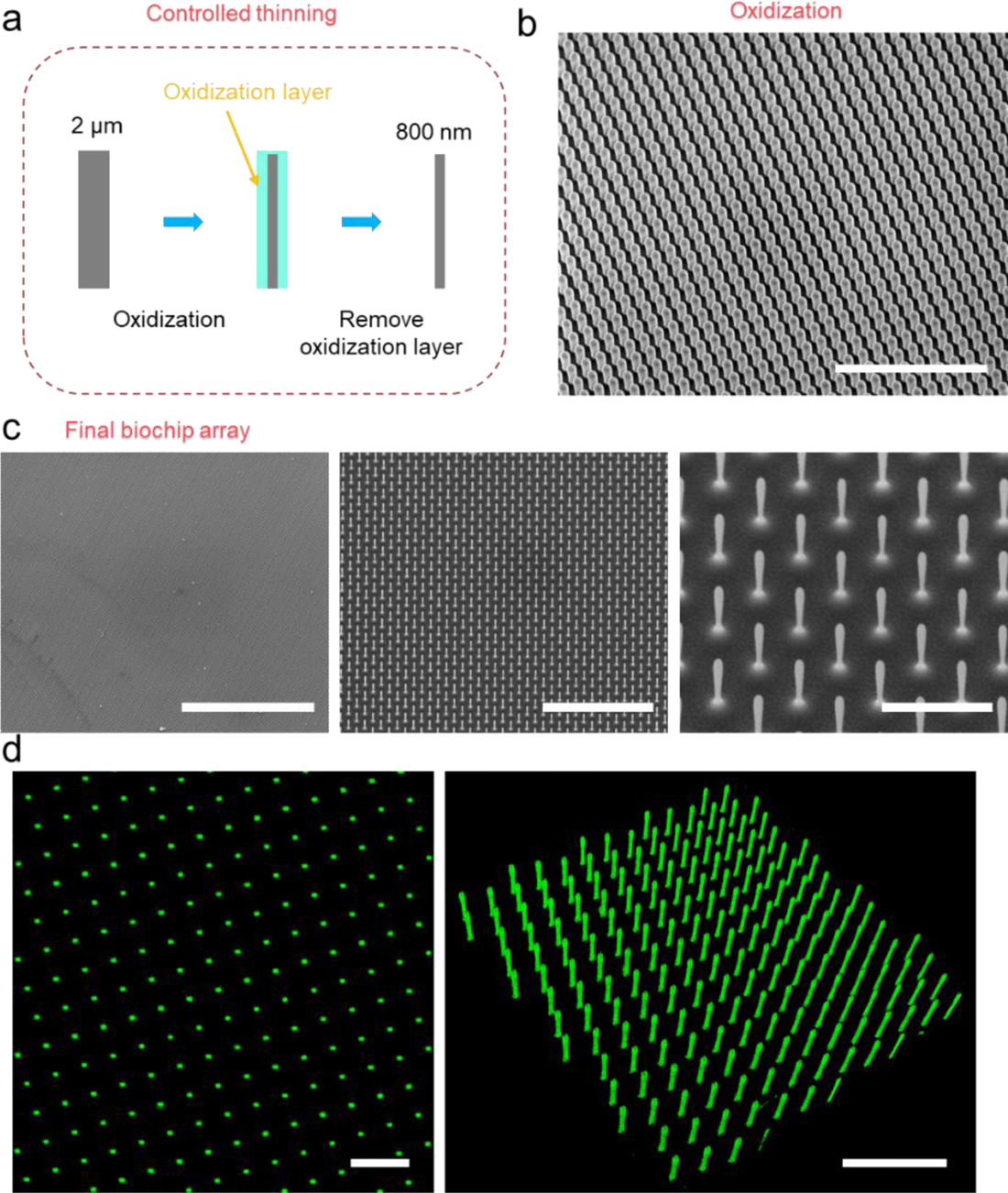
Fabrication of biochips with vertically aligned nanoprobes for post-transcriptional profiling. (a) Controlled thinning of silicon nanoprobes from a diameter of ∼2 μm to ∼800 nm. (b) SEM image of the oxidized silicon nanoprobes. Scalebar, 50 μm. (c) SEM images of the final nanoprobes at different scales. Scalebar, 400 μm (left), 50 μm (middle), 10 μm (right). (d) Representative fluorescence images of the nanonprobes, scale bar, 10 μm (left), 20 μm (right).

**Supplementary Figure S3.**
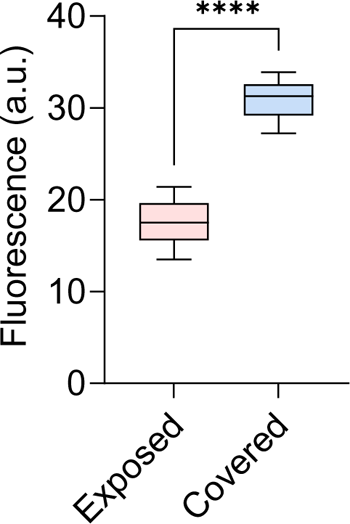
Generation of fluorescence contrast in nanoprobes with (high fluorescence) or without (low fluorescence) tissue coverage by UV-induced cleavage of fluorescence labels.

**Supplementary Figure S4.**
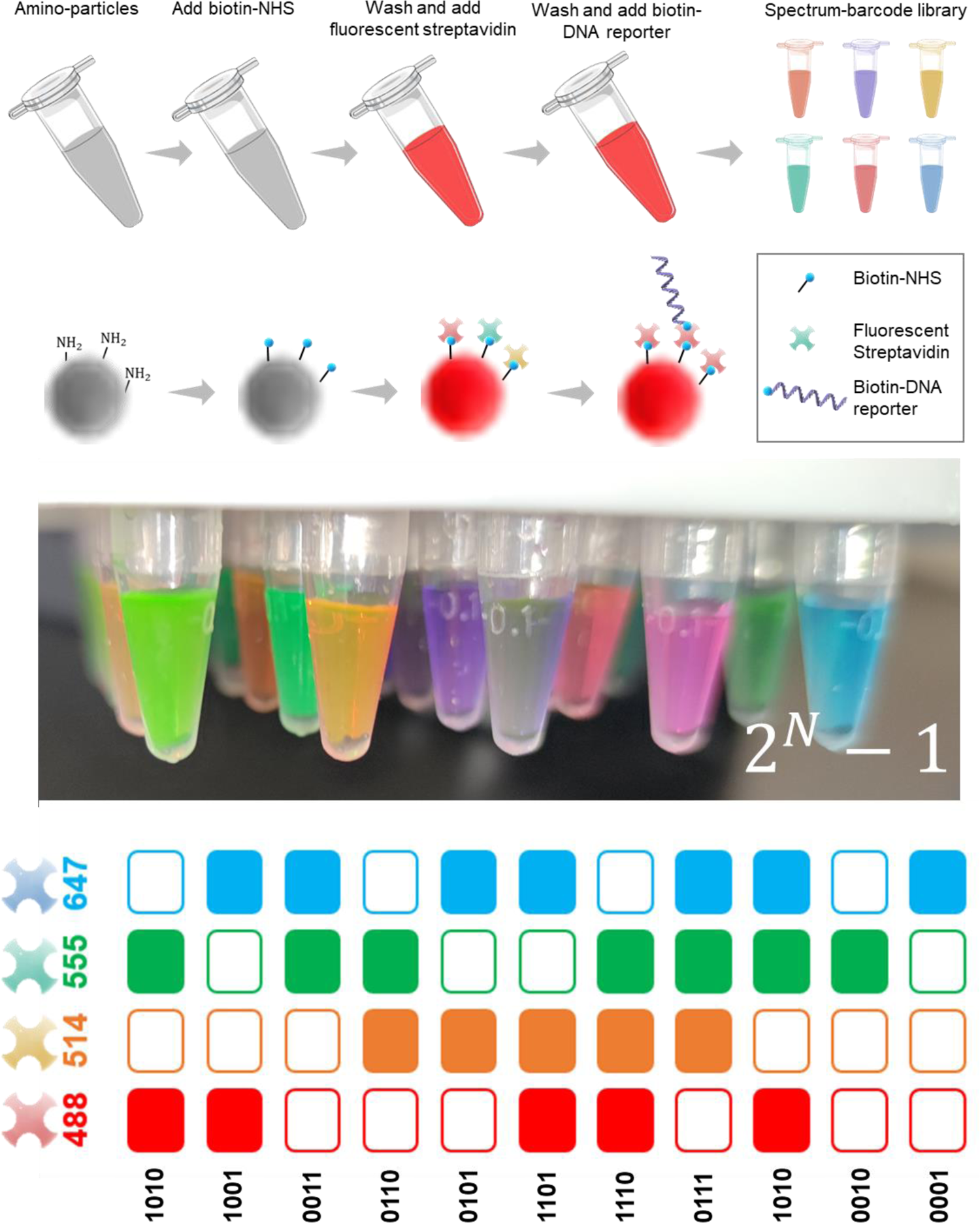
Schematic of Spectrum-CODE system.

**Supplementary Figure S5.**
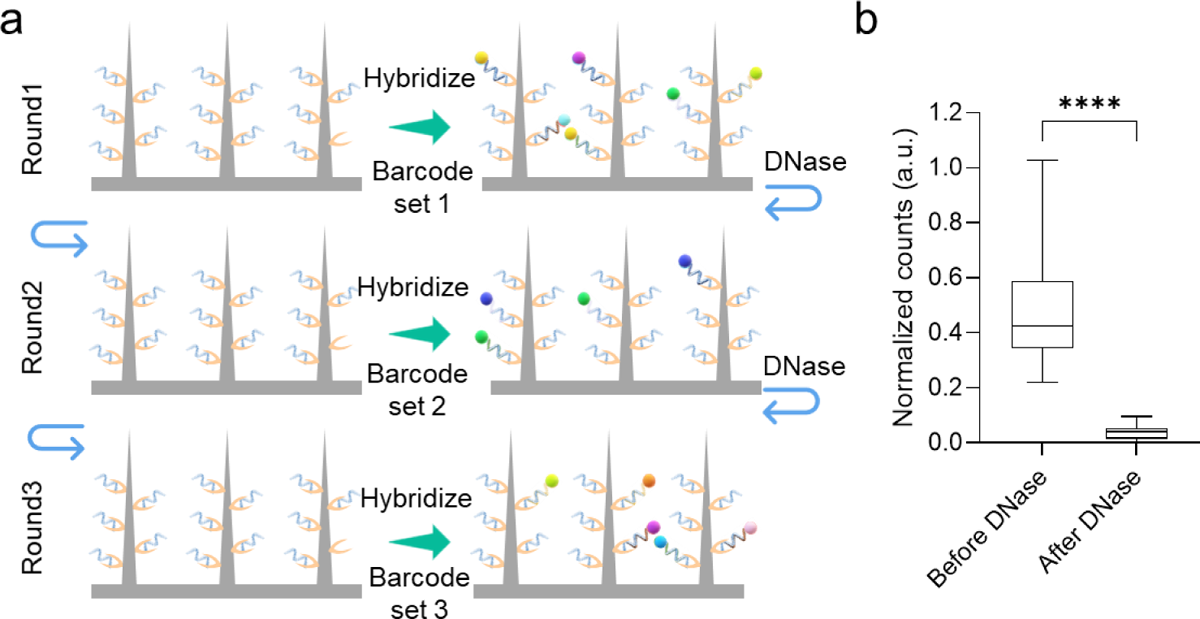
Multi-round hybridizations to increase the barcoding throughput. (a) Workflow of the procedures. (b) Comparison of bead counting before and after a DNase treatment.

**Supplementary Figure S6.**
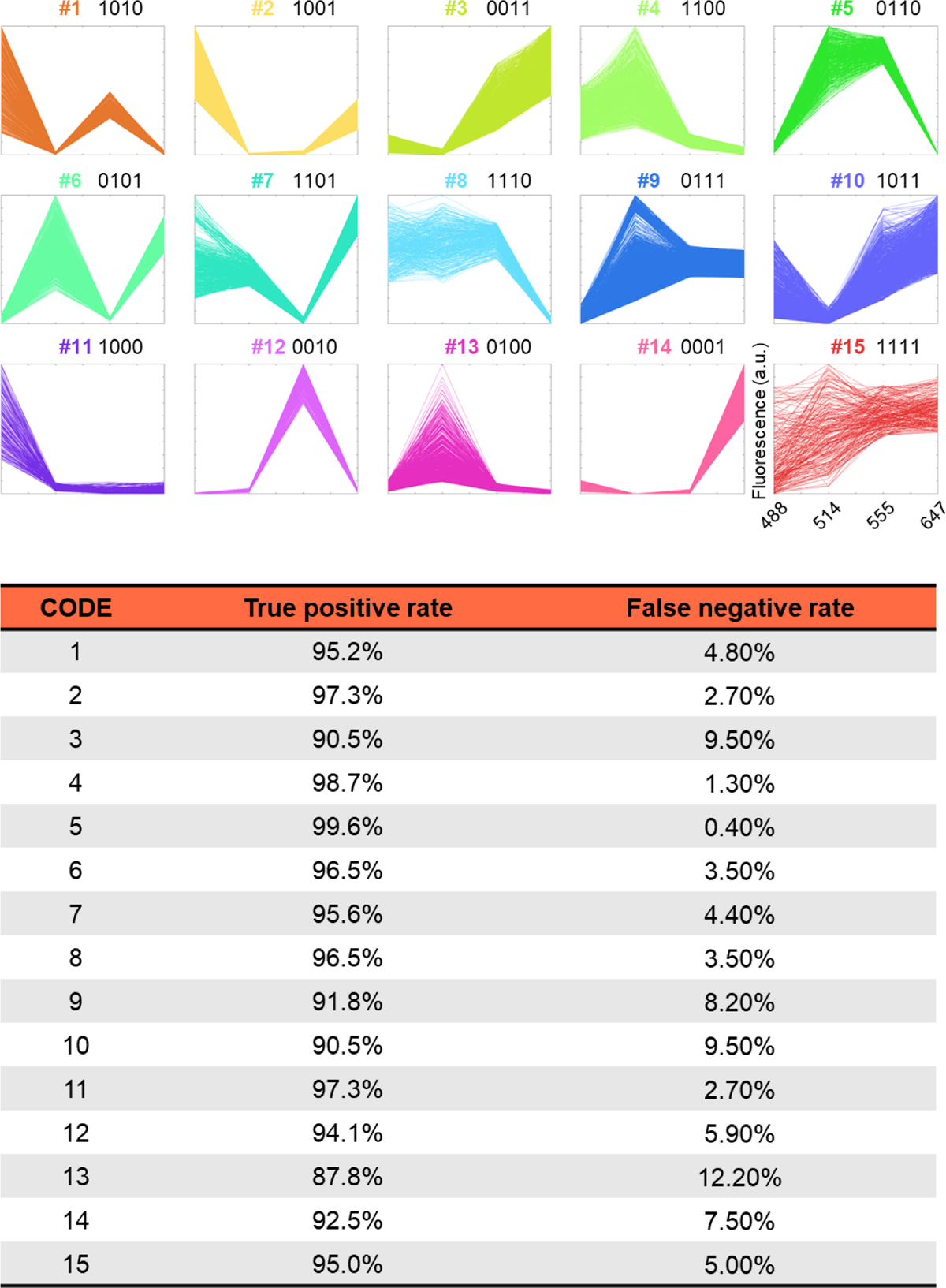
4-channel fluorescence parallel plots for nanobeads with 15 different Spectrum-CODEs and the true positive rate/false negative rate during machine learning recognition.

**Supplementary Figure S7.**
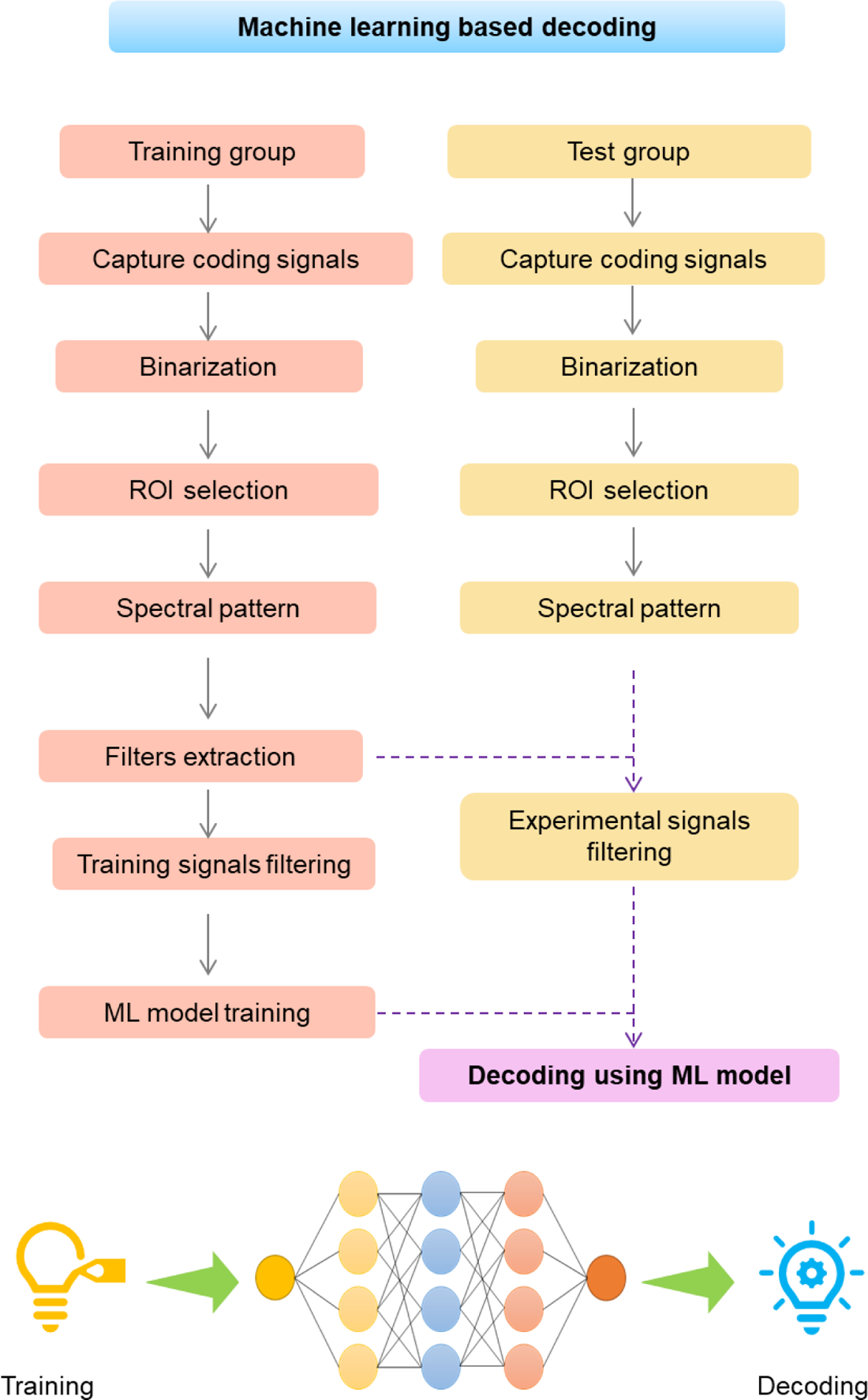
Machine leaning algorithms and workflow for spectrum recognition and multiplexed decoding.

**Supplementary Figure S8.**
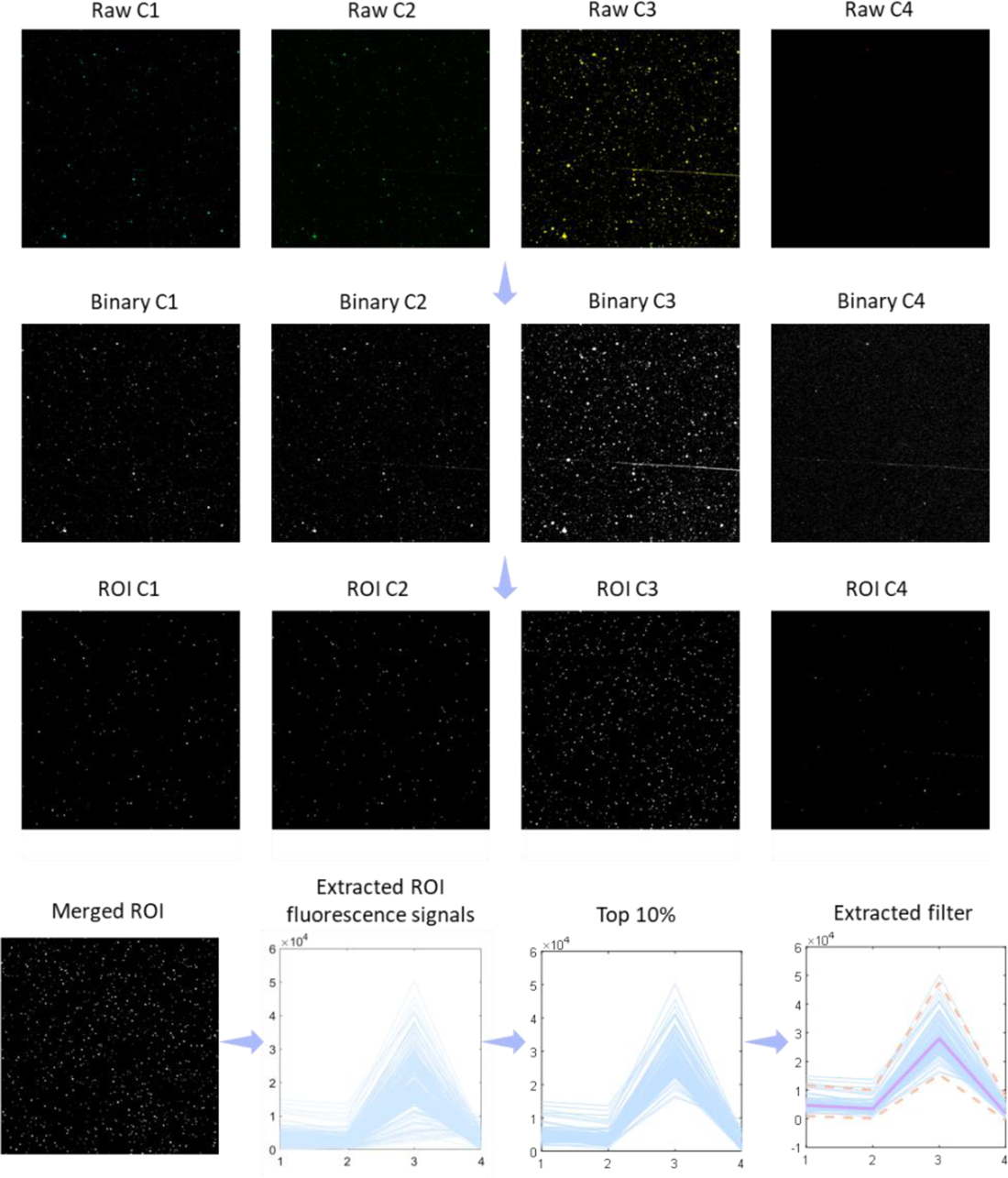
Image processing pipeline for ROI extraction and filtering based on the different fluorescence encoding.

**Supplementary Figure S9.**
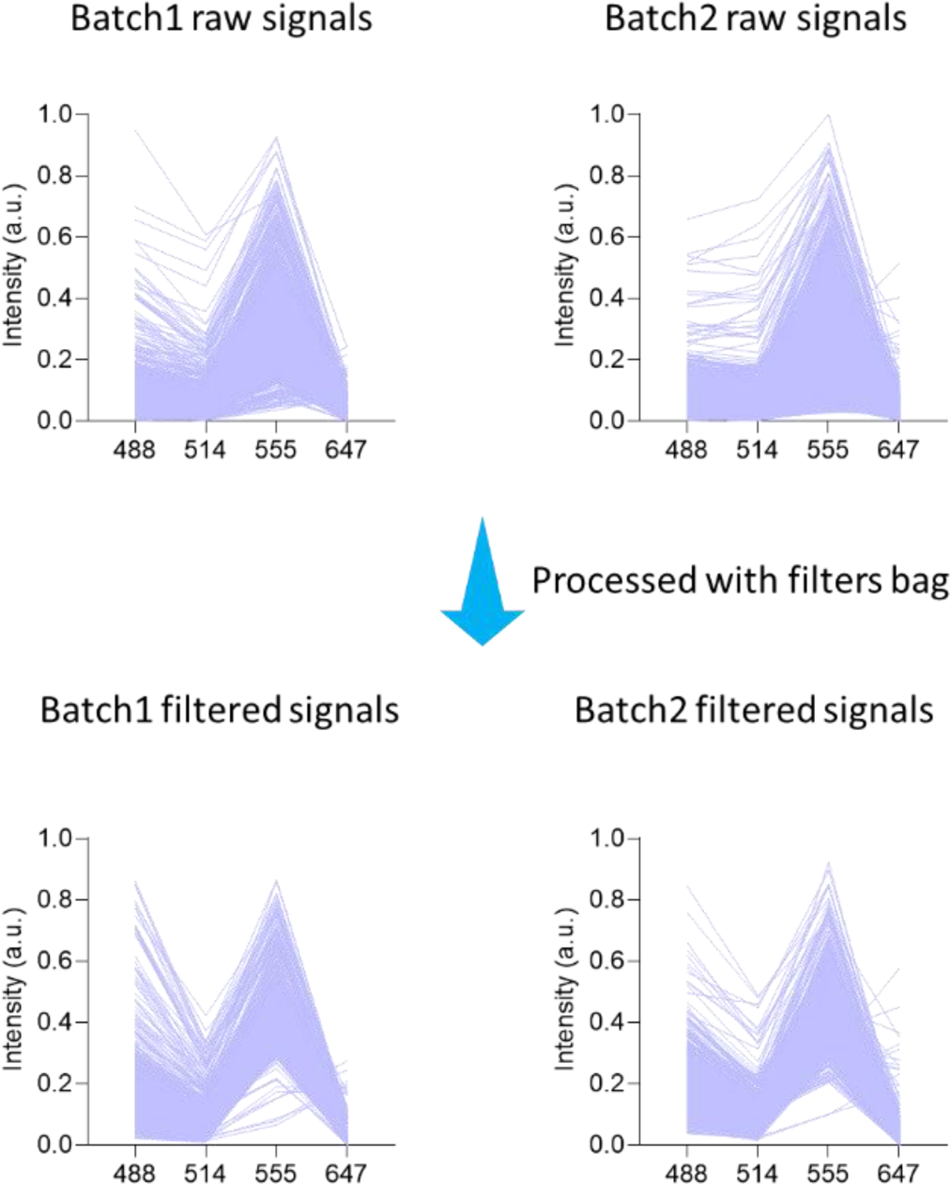
Analysis of batch variation for nanobeads with same barcode but were synthesized in different batches.

**Supplementary Figure S10.**
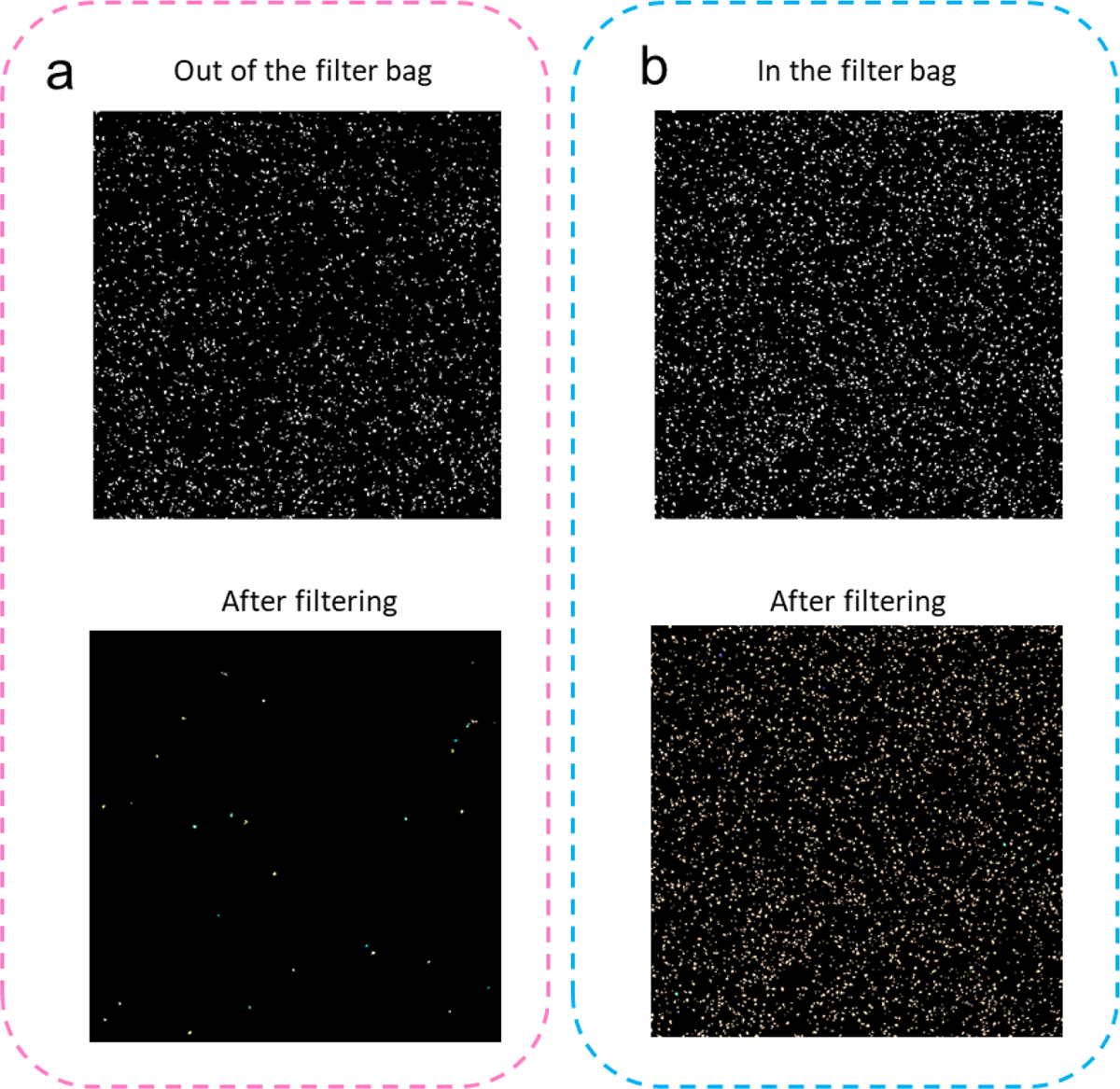
The filter bag system to remove microscopic noises. Barcodes out of the filters bag were removed after filtering (a), while barcodes within the filters bag are retained (b).

**Supplementary Figure S11.**
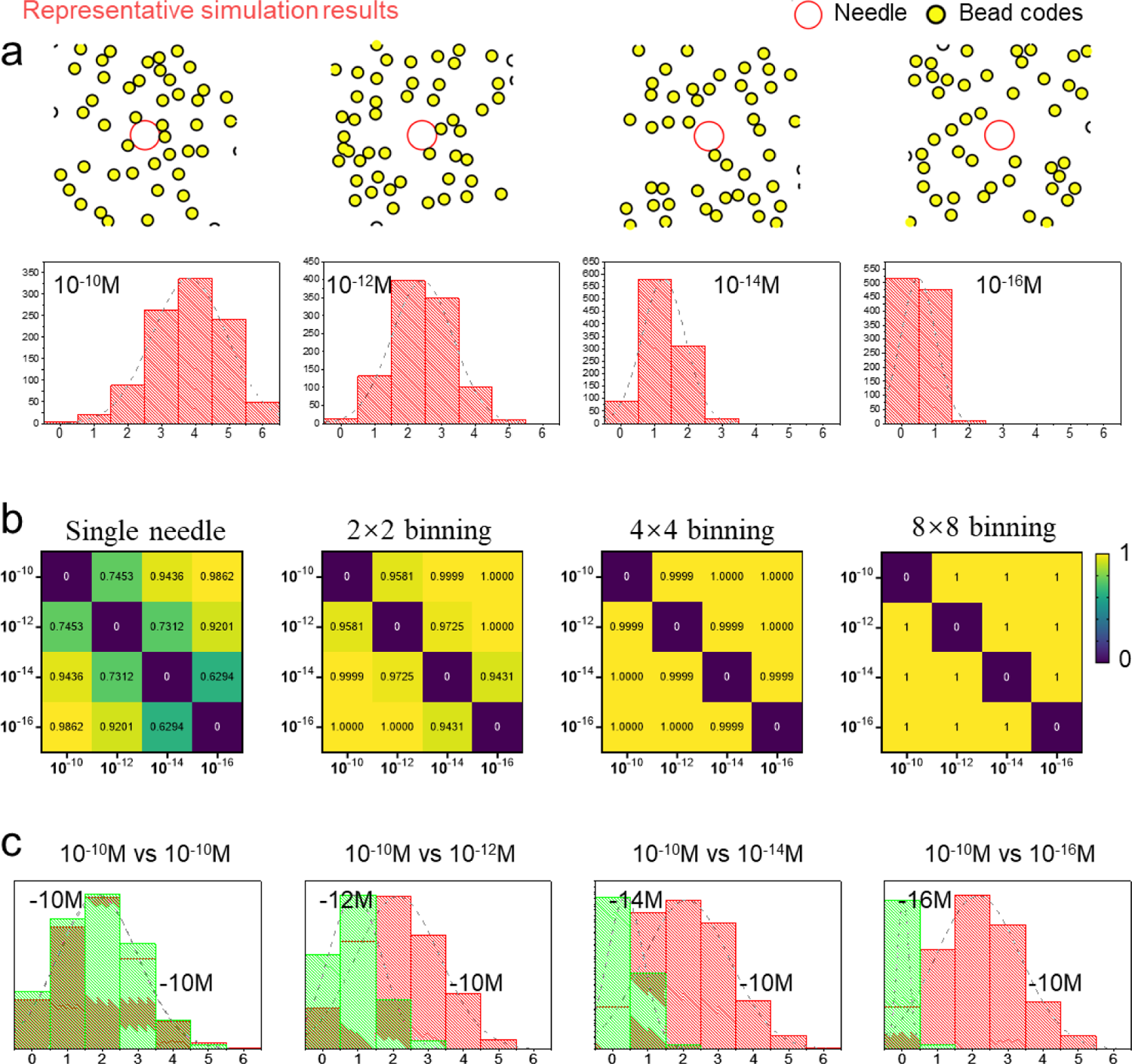
Probability based binding events in Spectrum-CODE beads system. (a) The binding events happen in a stochastic manner, where targets with higher surface intensity shows higher binding probability. A simulation was performed to demonstrate the relationship of the binding beads number and the binding probability. (b) After theoretical demonstration of the probability based binding events, a question was further raised that, when randomly compare binding numbers in different concentration condition, can we successfully differentiate them? Here, score matrix to indicate the differentiation accuracy in different binning condition were calculated. (c) What happened in multiplex condition when there is competition between different type of beads? To answer this question, the binding number distribution on the needle surface in two types of targets competition condition were simulated and summarized as histogram.

**Supplementary Figure S12.**
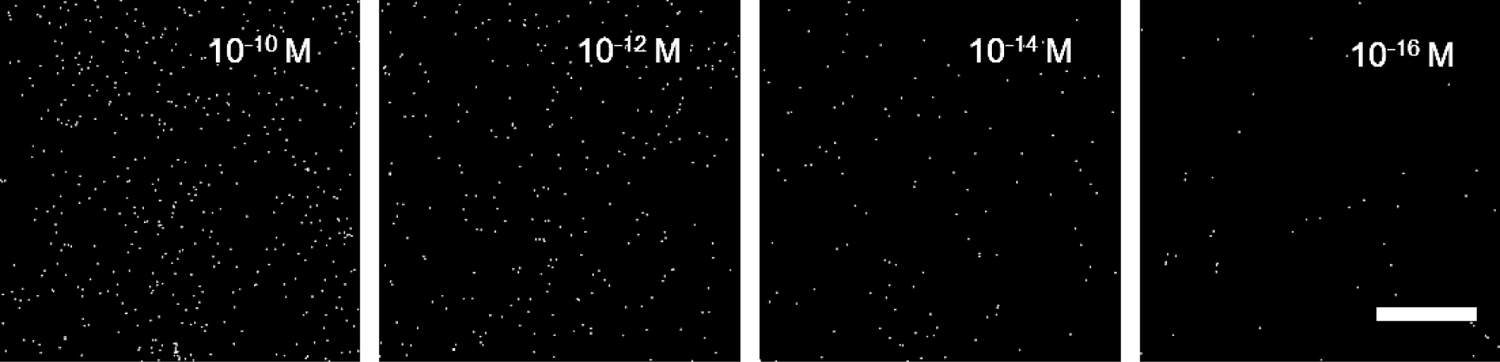
Spectrum-CODE beads binding on the biochip in different target concentration condition, Scale bar 50μm.

**Supplementary Figure S13.**
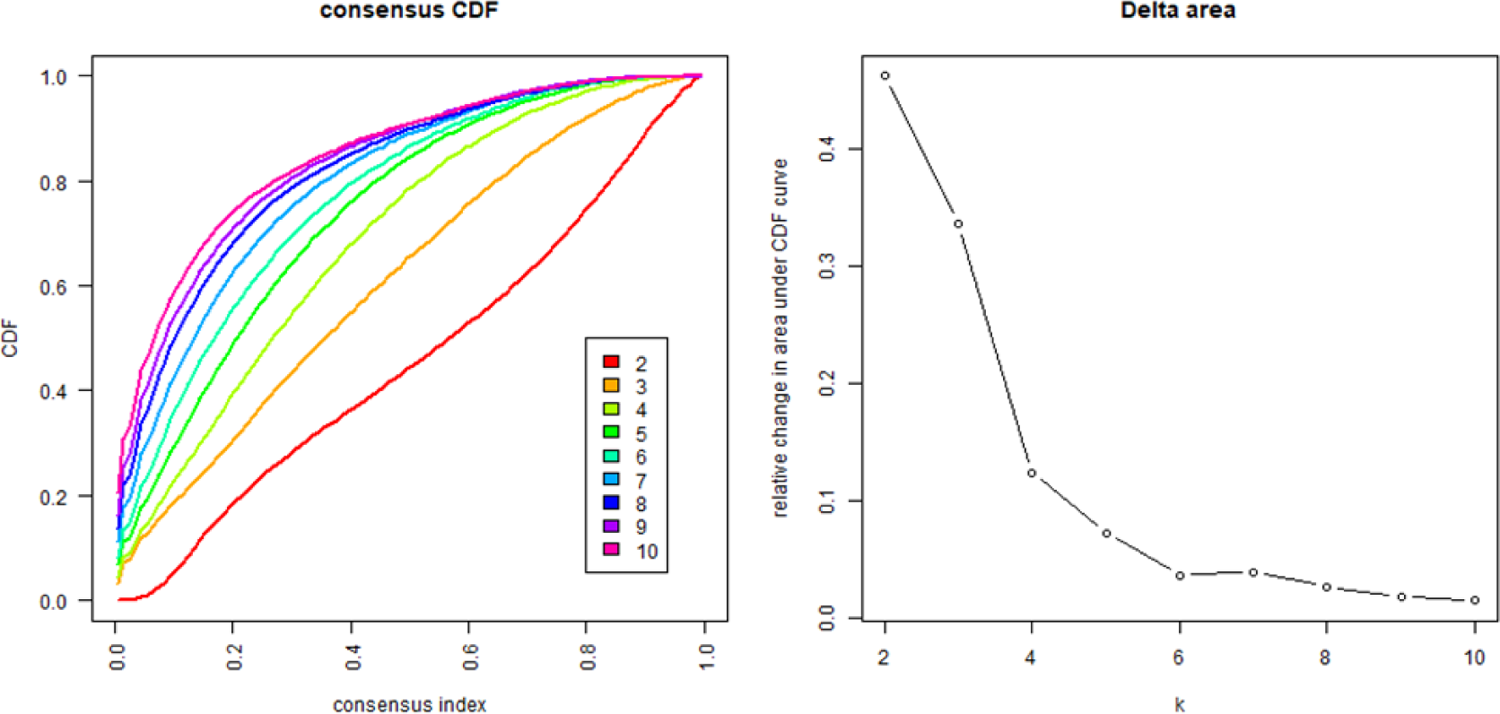
Consensus CDF and relative changes under CDF curve. The CDF plot allows a user to determine the relative increase in consensus and determine k at which there is no appreciable increase. The optimized cluster number (k=5) was selected based on the CDF delta area plot using the usual elbow method.

**Supplementary Figure S14.**
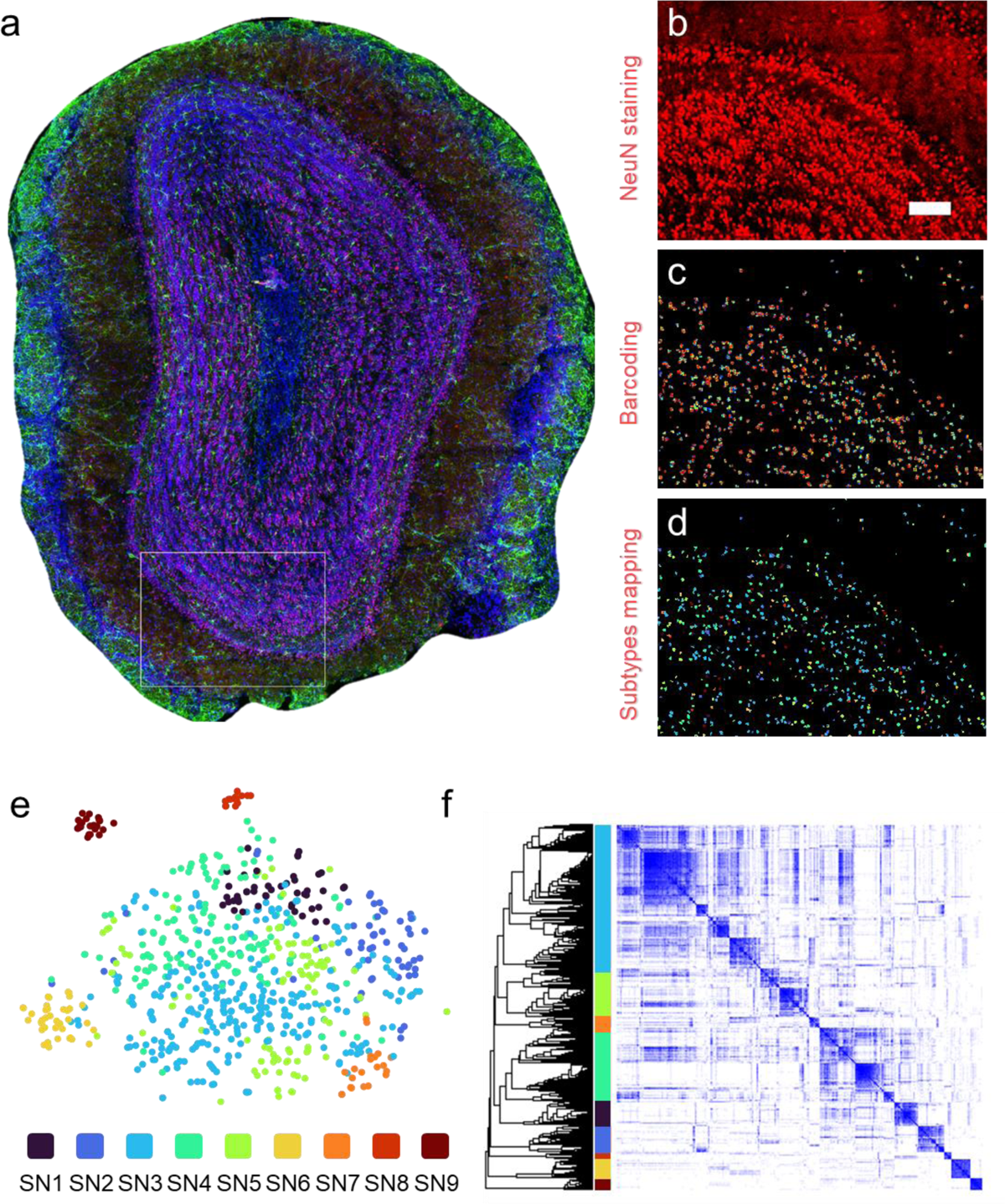
The heterogeneity of miRNAs for analyzing cellular subtypes in NeuN^+^ cells. (a) Immunostaining of the complete tissue. The rectangular region was used for the following analysis. (b) Immunostaining for NeuN to identify neuronal cells. Scalebar, 100 μm. (c) The nanoprobe-associated nanobeads with miRNA-specific spectrum codes, which are acquired by Spectrum-FISH analysis. (d) Spatial analysis of cellular subtypes in NeuN^+^ cells using unsupervised clustering of miRNA vectors. (e) The clustering of NeuN^+^ cells as shown in t-SNE distribution. (f) Clustering heatmap showing the single cell analysis of miRNAs expressions in NeuN^+^ cells.

**Supplementary Figure S15.**
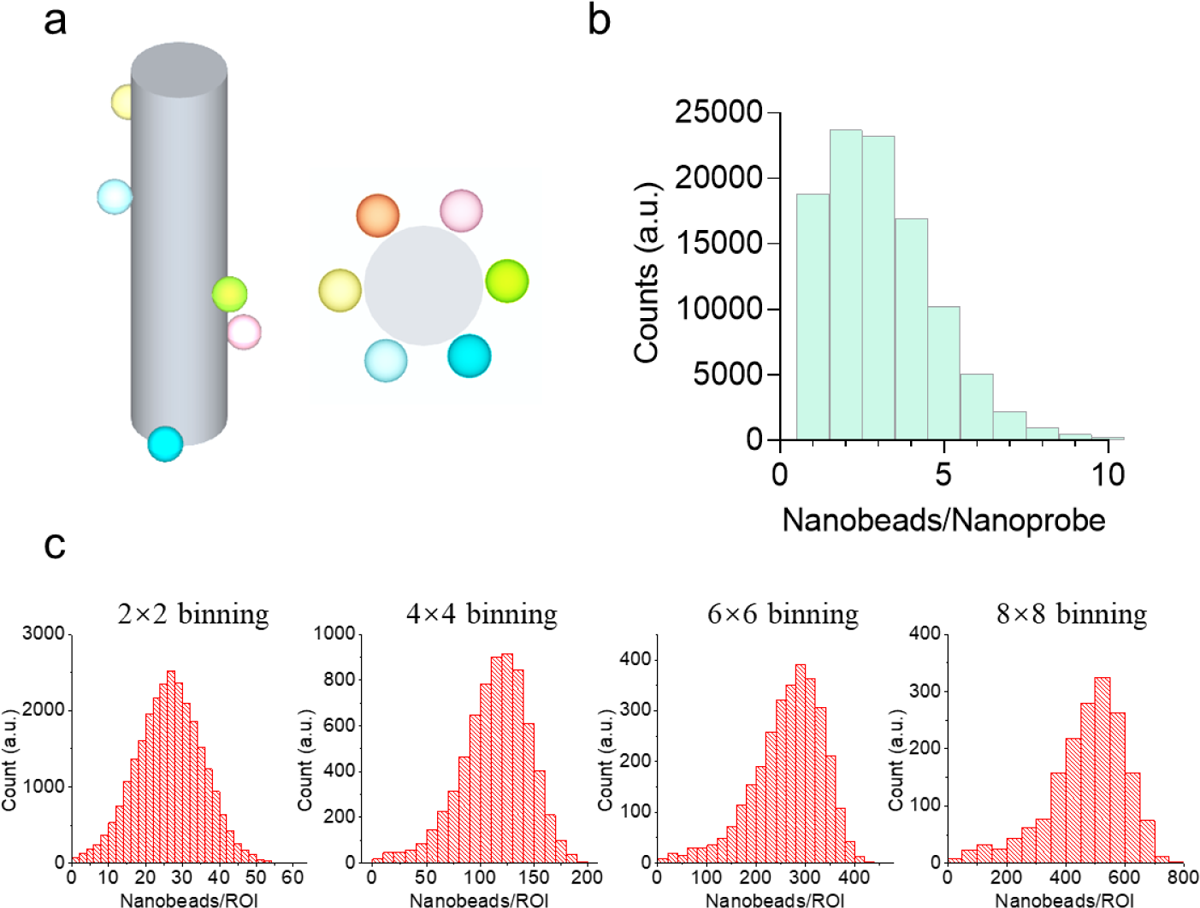
Analytical details of the spectrum codes acquired from the nanoprobes. **(a)** Theoretical evaluation of the binding condition of nanobeads (300 nm in diameter) on individual nanoprobes (800 nm in diameter). **(b)** Histogram of the nanobead counts on individual nanoprobes. **(c)** The effects of different binning on the quantification of nanobeads in the ROIs.

**Supplementary Figure S16.**
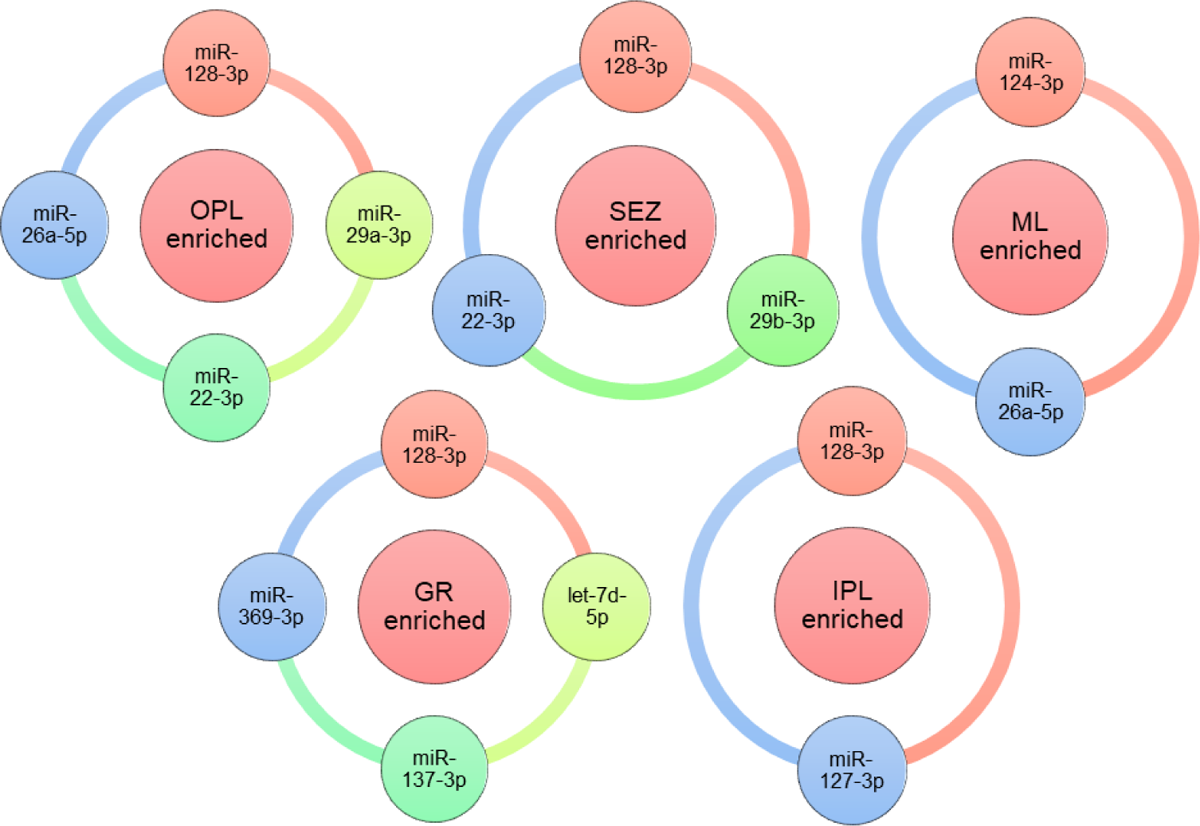
Visualization of the potential correlation between the miRNA signature for a cluster and its associated spatial enrichment in specific OB sub-regions.

**Supplementary Figure S17.**
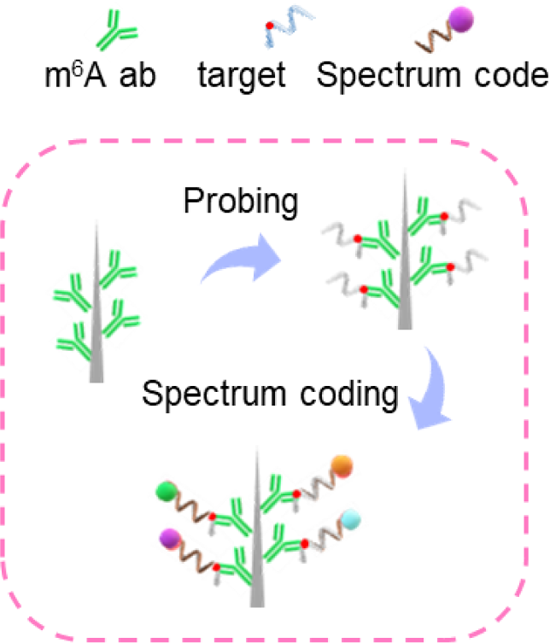
Nanoprobe functionalization strategy for molecular fishing of m^6^A-mRNAs using specific antibodies as the “bait” protein.

**Supplementary Figure S18.**
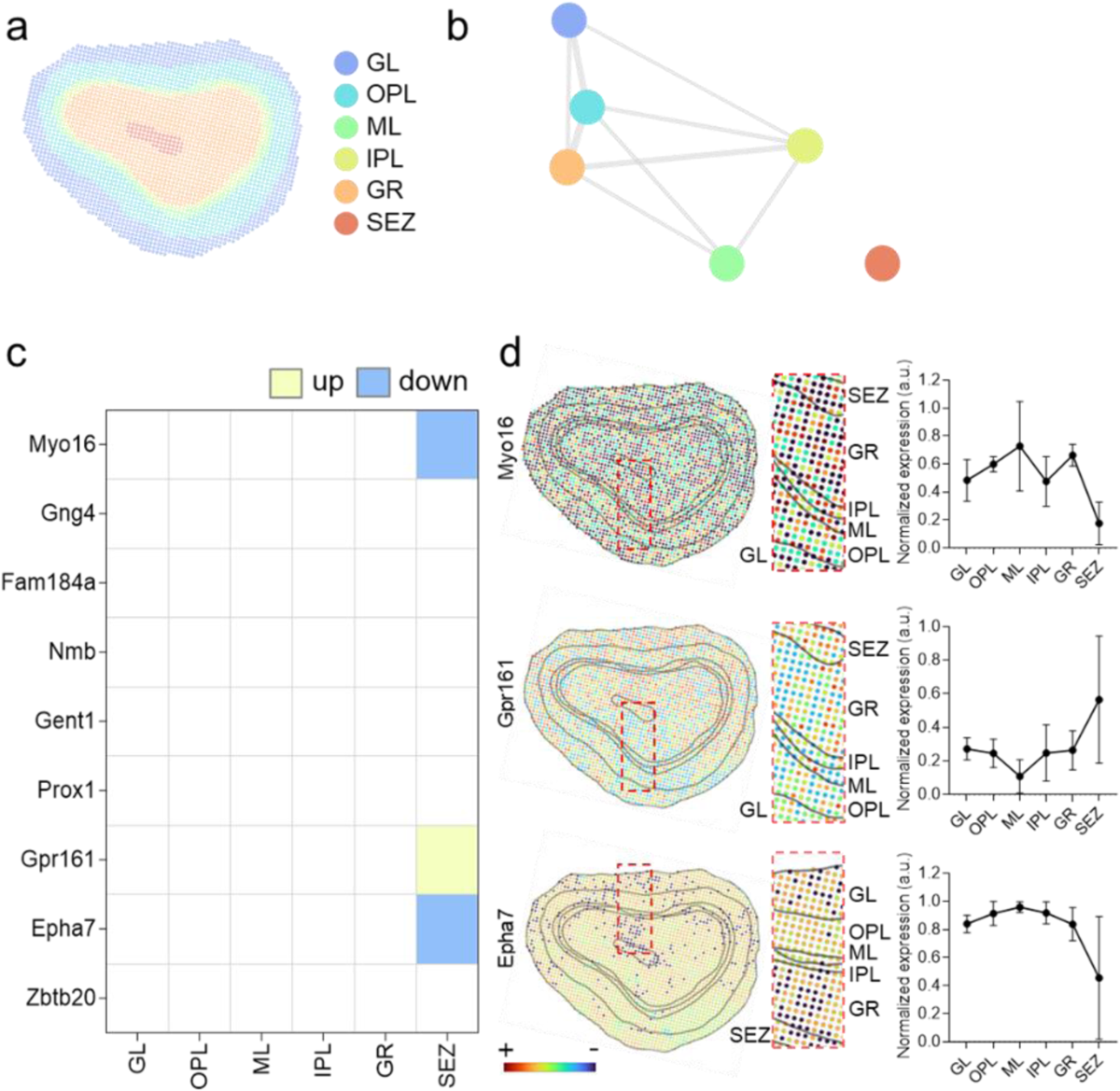
Regional organization of m^6^A-mRNAs in olfactory bulb. **(a)** Labeling of six anatomical regions in a coronal OB slice. **(b)** The similarity network of the six OB regions derived by using the associated m^6^A-mRNA data. Shorter distance and thicker connection represent higher similarity in m^6^A-mRNA expression. **(c)** Identification of the dominant m^6^A-mRNAs (out of the 9 targets) for each OB region. The comparison was made between one region versus all others, a significant target was defined as a fold change greater than 1.1 and a p<0.05 by unpaired t-test. **(d)** Representative spatial expression of a particular m^6^A-mRNA across a whole OB slice. The analysis was performed with a 4×4 binning of the nanoprobes, the error bars indicate mean ± SD, data from more than ∼200000 nanoprobes were collected from three biological replicates.

**Supplementary Figure S19.**
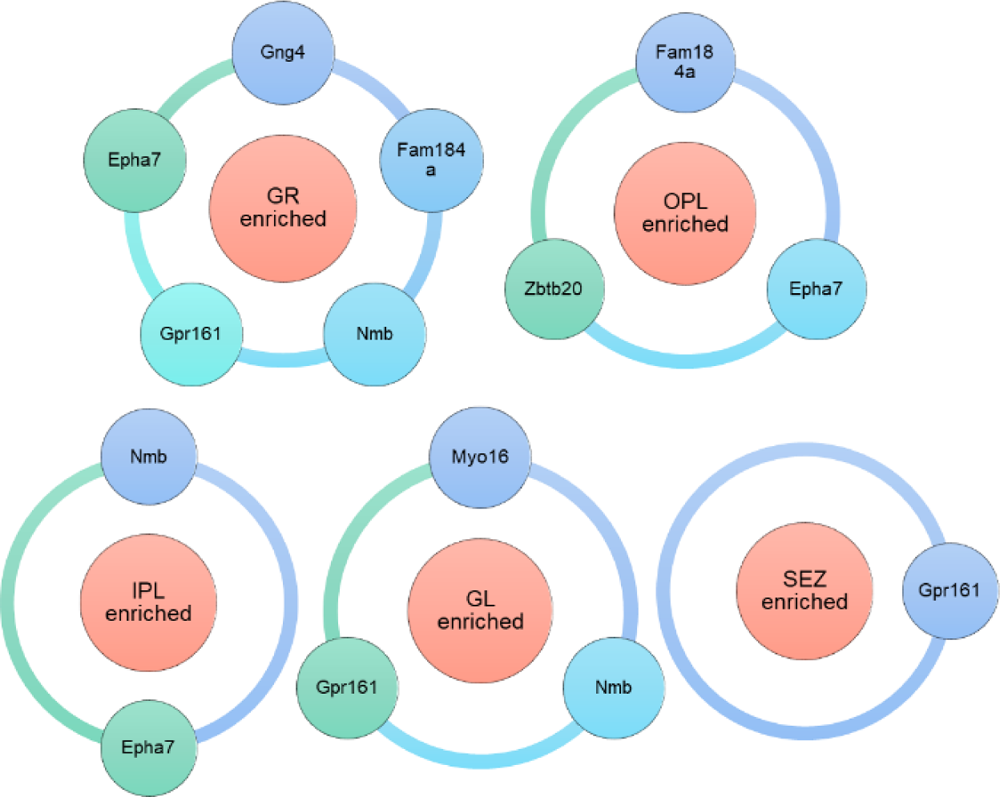
Visualization of the potential correlation between the m^6^A-mRNA signature for a cluster and its associated spatial enrichment in specific OB sub-regions.

**Supplementary Figure S20.**
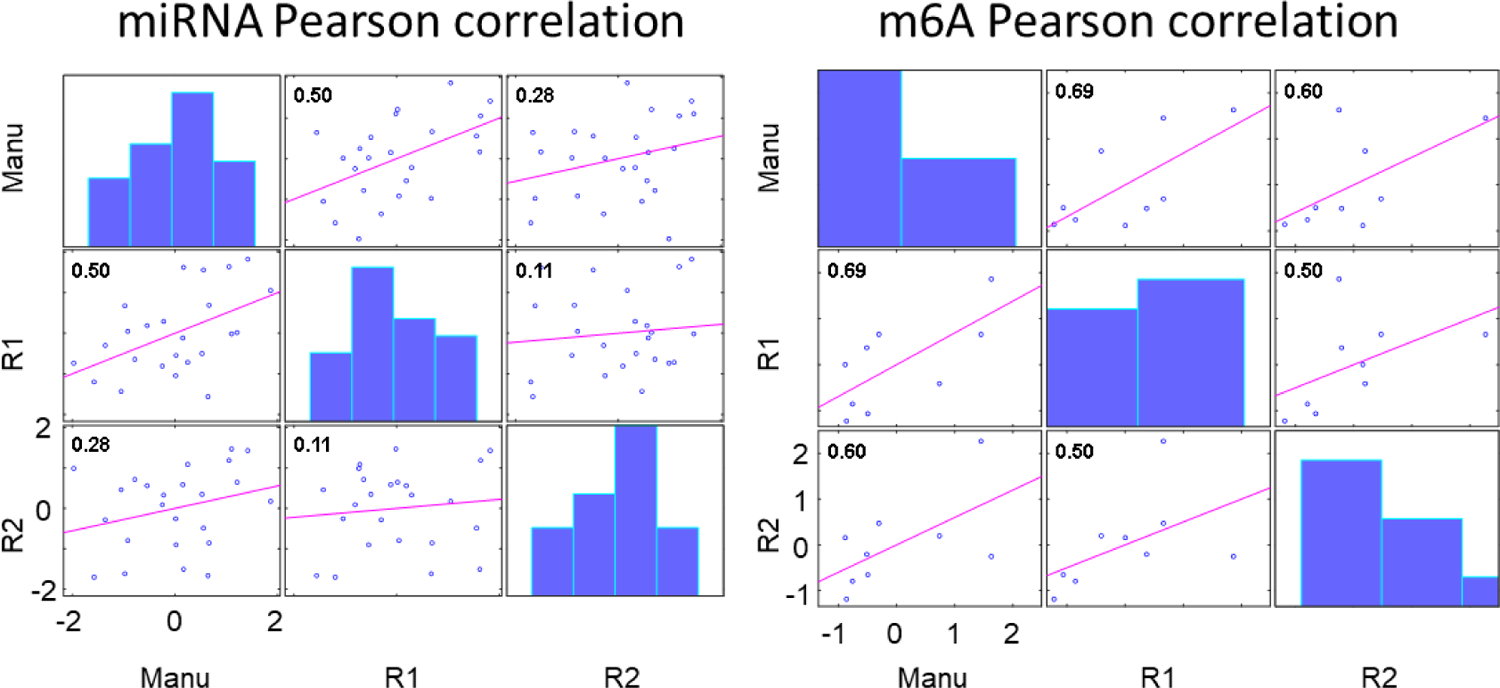
miRNA or m^6^A mRNAs expression Person correlation among replicative slices. ‘manu’ represents the data presented in the manuscript batch. ‘R1’ represents the replication 1 data batch. ‘R2’ represents the replication 2 data batch.

**Supplementary Figure S21.**
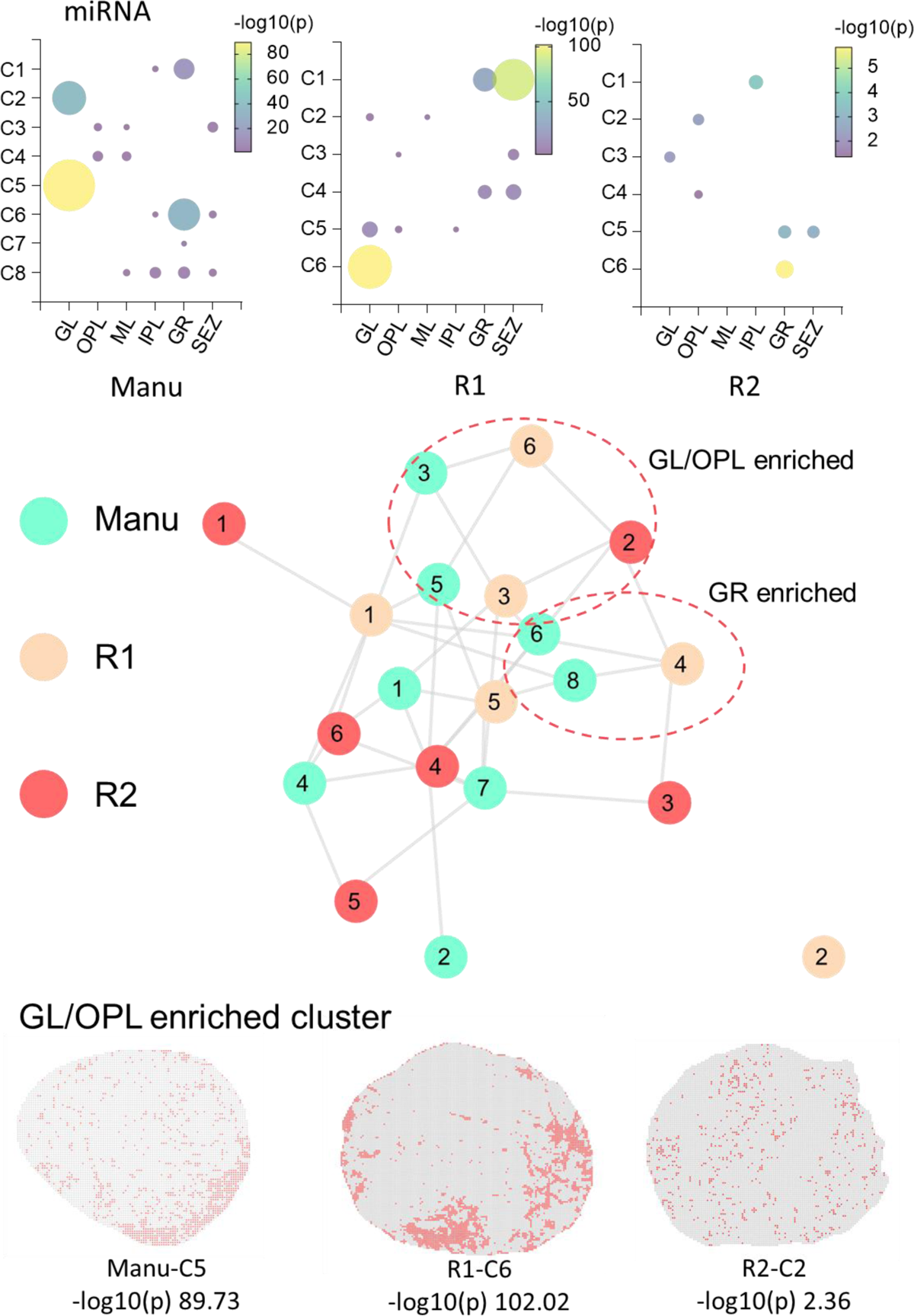
miRNA expression similarity between clusters in replicative slices. Hypergeometric test was performed to calculate the enrichment significance in particular OB regions (1^st^ row). An expression feature fingerprint was generated for each cluster and all the cluster pairs with Euclidean distance <0.4 were plotted on the graph with edge connection in miRNA cluster similarity network (2^nd^ row). A GL/OPL enriched region was found in the similarity network and representative GL/OPL enriched clusters with significance level were also presented (3^rd^ row). ‘manu’ represents the data presented in the manuscript batch. ‘R1’ represents the replication 1 data batch. ‘R2’ represents the replication 2 data batch.

**Supplementary Figure S22.**
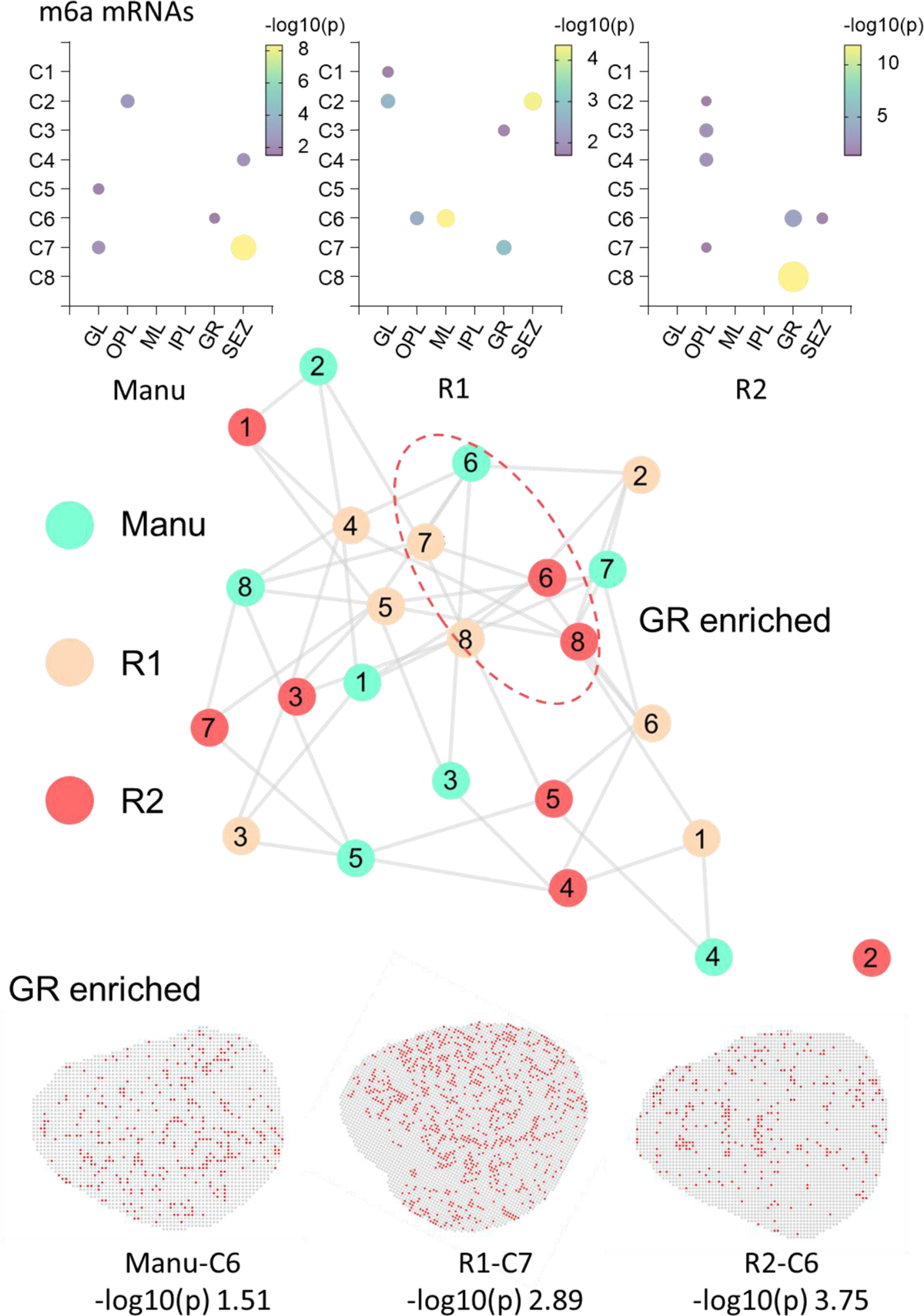
m^6^A-mRNA expression similarity between clusters in replicative slices. Hypergeometric test was performed to calculate the enrichment significance in particular OB regions (1^st^ row). An expression feature fingerprint was generated for each cluster and all the cluster pairs with Euclidean distance <0.4 were plotted on the graph with edge connection in m^6^A-mRNA cluster similarity network (2^nd^ row). A GR enriched region was found in the similarity network and representative GR enriched clusters with significance level were also presented (3^rd^ row). ‘manu’ represents the data presented in the manuscript batch. ‘R1’ represents the replication 1 data batch. ‘R2’ represents the replication 2 data batch.

### Note 1 Spectrum-FISH procedure

Here, we describe a Spectrum-FISH technique for multiplexed in situ spatial profiling of miRNAs or m^6^A-mRNAs in acute tissue slices. An array of vertical silicon nanoprobes were functionalized with ‘bait’ (in current setup, the ‘bait’ is RNA binding protein, p19 or m^6^A antibody) to directly pull multiple targeted miRNAs or m^6^A mRNAs out of cell cytoplasm in a few minutes. After the inCell-Biopsy operation, a UV imprinting procedure was performed to generate a pixelized tissue outline on the nanoprobe array. Then the tissue was removed from the biochip and each nanoprobe worked as a separated reaction plant for parallel in situ visualization and quantification of miRNAs or m^6^A mRNAs with Spectrum-CODEs. The detection limit can reach as low as 10^−14^ to 10^-16^ M (corresponding to ∼6000 copies/μL to ∼60 copies/μL), which is much lower than the abundance of even a single copy in a cell (considering the cell as a 20 μm diameter sphere, the concentration for single copy is ∼10^-13^ M). Using inCell-Biopsy, we demonstrated multiplexed tissue-wide spatial profiling of miRNAs or m^6^A mRNAs (**Supplementary Figure S1**).

### Biochip functionalization

In the beginning of Spectrum-FISH technique, based on different spatial profiling purpose, the biochip can be specifically functionalized targeting on various type of molecules. such as p19 protein for miRNAs or antibody for m^6^A-mRNAs, as demonstrated in this study. Such a versatility can be readily extended to other ‘baits’, such as poly(T) sequences for mRNAs, different antibodies for signaling proteins or methylated RNAs, even RNA-binding proteins (RBPs) for interactive translational regulation factors. Multi-omics spatial epi-transcriptome analysis can also be explored by a “dual-bait” functionalization of the nanoprobes, so that the interactive roles of miRNAs and specific methylated mRNAs can be further explored with more delicate specificity and spatial dynamics, which could potentially provide new insights to the progress and treatment of human diseases. The biochip functionalization procedure mainly includes Piranha and (3-Aminopropyl)triethoxysilane (APTES) treatment in sequence to activate the nanoprobe surface with amino group, followed by reacting with biotin-NHS to further functionalize the nanoprobe surface with biotin. Finally, streptavidin (SA) was used to bind ‘baits’ molecules with biotin tag on the nanoprobe and finish biochip preparation.

### Molecular fishing

The inCell-Biopsy technique is based on the continuous development of a “molecular fishing” system, which uses an array of silicon nanoprobes as fishing rods for minimum-invasive access of cytoplasmic regions of mammalian cells. Specifically, for “fishing” miRNAs, a size-dependent RNA binding protein, p19, was cross-linked to functionalize the nanoprobes, working as the “fishing hook” to capture double-strand RNAs (dsRNAs). P19 can selectively bind to all dsRNAs of 20 to 22 base pairs (bp), which is a range covering almost all miRNAs in mammalian cells. When the prepared silicon nanoprobes are interfaced with cells using a centrifugation facilitated procedure, the fishing rods can penetrate the cell membrane to access the cytoplasmic region via a temporary membrane disruption. Upon the retrieval of the nanoprobes, the targeted miRNAs are isolated.

### UV imprinting

In the Spectrum-FISH system, just after the inCell-Biopsy procedure when the tissue was still in contact with the biochip, the spatial encoding was facilitated by a UV imprinting process, which gives a snapshot of the tissue morphology and boundary on the nanoprobe biochip and provides an associative reference linking the cell and nanoprobe spatial coordinate. The nanoprobes were labelled with fluorophores by photocleavable crosslinker. When a piece of tissue slice (smaller than the chip) was interfaced with the nanoprobes for “molecular fishing”, a brief UV (365 nm, ∼5 mw/cm^2^) irradiation was applied. This energy level was just enough to cleave off the fluorescent labels on the nanoprobes in direct UV exposure, while the rest part covered by the tissue remained unaffected, generating a pixelated fluorescent pattern of the tissue, which can be used to register of the nanoprobes to individual cells at later analytical stages.

### In situ analysis via Spectrum-CODE

After the UV imprinting procedure, the silicon nanoprobes biochip was separated from the tissue slice with the target molecules extracted by the ‘bait’. For multiplex visualization and quantification of the captured biotargets, we developed a digital spectrum method (Spectrum-CODE) for functional encoding of the analytical targets. The Spectrum-CODE can overcome the challenge of limited numbers of fluorescent channels and increase the multiplexing throughput of the Spectrum-FISH technique. The extracted biomarker molecules (e.g. miRNA or m^6^A-mRNA) were visualized by an in-situ hybridization with a “reporter” DNA, which was specifically designed to recognize and encode different targets. Each ‘reporter’ sequence was pre-labelled by amino-Fe3O4 nanobeads with a rainbow fluorescence composition. The simplest spectrum digitization method uses a “1” (with dye) or “0” (without dye) mixing to compose the spectral codes. Specifically, for analyzing RNAs, a DNase-assisted multi-round visualization process further expand the code pool. As demonstrated here, we used a four-channel system to encode 27 miRNAs, and the machine learning based spectrum decoding gives more than 90% accuracy. Theoretically, the encoding pool can be further expanded if the rainbow mixing changes from the current binary format and adopts a stepwise ratio for different fluorophores. In a trinary system with 7 different fluorophores would substantially increase the multiplexing throughput to over 10,000 by 5 rounds of visualization. It is worth mentioning that this over 10000 throughput is the theoretical limit of the Spectrum-CODE system in specific condition.

### Note 2

#### Spectrum-CODE system for molecular sensing

The Spectrum-CODE system for molecular sensing includes high-throughput individually labelled beads with unique fluorescence code, which were further decoded by machine learning algorisms for quantification. By functionalization of specific antisense oligoes or antibodies on the surface of the encoded fluorescent beads, followed with specifically binding or hybridization with protein (achieved by antibodies) or DNA, RNA (achieved by antisense oligoes), the molecule number of related protein or DNA, RNA can be counted by counting the number of the encoded fluorescent beads. Generally, in this way, we can achieve high throughput detection, identification and quantification of DNA, RNA, and protein targets.

#### Materials aspect

For production of unique fluorescent encoded particles. In material aspects, the fluorophores used for coding can be common commercial fluorescent molecules, such as FAM, Cy3, GFP proteins, and Alexa fluor series provided by thermoFisher (Alexa fluor 488, alexa fluor 555,.etc), quantum dot and up-conversion nanoparticles and any other materials that can emit fluorescence. The shape of the encoded particles can be spherical, nanorod, nanowire, star and any other shapes. The materials of the encoded particles can be general magnetic materials, such as Fe_2_O_3_, Fe_3_O_4_ and can also be silicon, SiO_2_, gold, Ag and AlOOH. The materials can also be various polymers materials such as polystyrene and polyvinyl chloride.

#### Spectrum-CODE encoding principle

In fluorescence coding manner aspects, the way to apply the fluorescence coding on the beads is through binding single bead with fluorophores in different combinations. For example, if we have three types of fluorophores, namely fluorophore A, fluorophore B and fluorophore C. The existence of each fluorescence can be indicated by two statuses, namely ‘1’ and ‘0’, where ‘1’ means existence and ‘0’ means no existence of related fluorophore. Therefore, if this single particle simultaneous have fluorophore A, B, C, we can write this fluorescence digital code as ‘1 1 1’, where the first ‘1’ represents the existence of fluorophore A, the second ‘1’ represents the existence of fluorophore B and the third ‘1’ represents the existence of fluorophore C. Similarly, if we got another single particle only have fluorophore A and B, then the fluorophore digital code is ‘1 1 0’. Here we can then finally conclude that with three types of fluorophores (N=3), we can have in total 7 fluorophores digital codes, listed as ‘1 1 1’, ‘1 1 0’, ‘1 0 1’, ‘0 1 1’, ‘1 0 0’, ‘0 1 0’ and ‘0 0 1’. The ‘0 0 0’ is not used since it appears no fluorescence. We can further extend this scenario into more fluorophores with the following equations, 2^N-1, where N is the number of fluorophores used for coding.

Further, in above calculation, we only consider two fluorescent statuses, namely ‘1’ and ‘0’, where ‘1’ means the existence of the fluorophore or can be further indicated by ‘bright’ and ‘0’ can be indicated by ‘dark’. Here we can extend these two fluorescence statuses (‘bright’ and ‘dark’) for single fluorophore to more fluorescence statuses by the introduce of ‘fluorescence intensity’. For example, we can have three fluorescence statuses, namely ‘0’, ‘1’, ‘2’ in digital code and representing ‘dark’, ‘half-bright’, ‘bright’ fluorescent statuses, respectively. Therefore, in three fluorophore condition (N=3) and three fluorescence statuses (I=3), we will get in total 26 fluorescence codes. The equation can be I^N-1, where I indicates the number of fluorescent statuses caused by fluorescence intensity, N indicates the number of fluorophores used for coding.

#### Crosslinking methods

Based on above description, the particles will show different fluorescent combinations by binding the beads with fluorophores. The binding process can be achieved in several widely used methods. a) Electrostatic adhesion. In this condition, the particles surface will carry opposite charges with that of fluorophores. For example, if the fluorophores have negative surface charge in solution and the particles have positive charges. Then they will tend to bind with each other by the electrostatic adhesion. b) Absorption. If the particles are porous, then the fluorophores can be absorbed into the inside pores to achieve the binding status between particles and fluorophores. c) Crosslinking. The fluorophores and the particles can be bound together by crosslinking process. The crosslinking process can be different in various conditions. i) fluorophores and particle both have amino group and can be crosslinked via glutaraldehyde or NHS-PEG-NHS; ii) fluorophores and particles can be crosslinked via biotin-streptavidin. iii) fluorophores and particles can be crosslinked via NHS-NH_2_ reaction, for example, Alexa-fluor 488-NHS with amino-Fe_3_O_4_ particles. The fluorescence signals are recorded and imaged by fluorescence microscope.

#### Machine learning algorism for decoding

For decoding (readout and identification of the fluorescence codes), it is hard for us to directly identify them based on visual observation or determination by threshold because of the overlapping of the spectrums emitted by the fluorophores. This cross-channel interference will lead to the existence of weak fluorescence in some fluorescent channel, where the corresponding fluorophore doesn’t really exist. Besides, the differentiation of various fluorescence intensity makes this identification process even more difficult. Therefore, we used common machine learning schemes to assist this readout and identification process. To be stated, in this part, generally most existed machine algorisms can be used. The difference might be the accuracy, some might be better and more suitable for decoding these fluorescent codes. We can use traditional machine learning algorisms such as linear regression, logistic regression, decision tree, SVM, Bayes and KNN, etc. We can also use the advanced deep-learning algorisms, such as Convolutional Neural Networks (CNNs), Long Short Term Memory Networks (LSTMs), Recurrent Neural Networks (RNNs), etc. During applying these different algorisms, we will for starter prepare a training groups dataset, including the actual fluorescence intensity distribution of all the fluorescence codes used. Then we will obtain a pretrained machine learning model. For decoding, an unknown fluorescent encoded particle will be imaged and recorded. The recorded fluorescence signals will be input in the pretrained machine learning model and then be decoded to obtain its identity in a format of the digital code.

### Note 3

#### Probability based binding events in spectrum-CODE beads system

After ‘molecular fishing’, although there are thousands of targets on the needle surface, the beads won’t bind all the targets due to the stereo-hindrance effect. The binding number is an indicative value corresponding to the target intensity on the needle surface. The binding events happen in a stochastic manner, where targets with higher surface intensity shows higher binding probability. A simulation was performed to demonstrate the relationship of the binding beads number and the binding probability.

Here, beads with Brownian motion were used for simulation, where all the beads have a random birth location, random motion direction and random motion distance in each simulation step. The beads were all assigned with 300 nm diameter and an 800 nm needle with red outline was placed in the middle of the simulation area. When a bead randomly contacts with the needle surface, the bead will be frozen and simulating the binding events. This frozen action was assigned to happen with different probability (The 10^-10^ M was set as 1/10 binding probability, 10^-12^ M, 1/20 probability, 10^-14^ M, 1/40 probability, 10^-16^ M, 1/80 probability), where the binding probability were all defined according to the real binding number distribution in experimental condition (**Supplementary Figure S15**) and try to indicate the real status. It is worth to note that the intrinsic nature of the relationship between 10^-10^ M and the different binding probability could be summarized as follows: target concentration in solution influences target absorbance intensity on needle surface and the target intensity on needle surface will lead to different binding probability of the corresponding target specific beads. For example, in a 10^-12^ M condition, each time the beads contact with the needle surface, the probability of the binding events happening was 1/20, which means that almost every 20 contact events happen, there will be in average one binding event. The simulation was set as in total 100 steps, where all the beads have 100 times random motion during the whole simulation (**Supplementary Figure S11**).

After each time simulation, the final motion frame shows the binding status. A series of representative simulation results were presented (**Supplementary Figure S11a**). For example, in **Supplementary Figure S11a**, the final binding numbers were 4, 2, 1, 0 in 10^-10^ M, 10^-12^ M, 10^-14^ M and 10^-16^ M condition, respectively. Notably, only single time simulation will have too much randomness. Therefore, we performed 1000 times simulations and record the results (**Supplementary Figure S11b**), where the 10^-10^ M shows a binding number distribution range from 0 to 6 with a maximum point at 4, the 10^-12^ M shows a binding number distribution range from 0 to 5 with a maximum point at 2, the 10^-14^ M shows a binding number distribution range from 0 to 3 with a maximum point at 1, the 10^-16^ M shows a binding number distribution range from 0 to 2 with a maximum point at 0. The simulation results clearly show that probability based binding events can have distinguishable difference in different concentration condition. Therefore, in current setup with stereo-hindrance effect where the maximum binding number in single needle slice section is 11 can still indicate the concentration difference.

#### When randomly compare binding numbers in different concentration condition, can we successfully differentiate them?

After theoretical demonstration of the probability based binding events, a question was further raised that, when randomly compare binding numbers in different concentration condition, can we successfully differentiate them? To answer this question, we prepare 1000 rounds comparison. In each round, we randomly select one binding result in each concentration condition and compare them, where if the binding number in higher concentration condition was larger than the lower concentration condition, the value was recorded as ‘1’ and if not, the value was recorded as ‘0’. After 1000 rounds random comparison, we generate a score matrix to indicate the differentiation accuracy. To further improve the robustness of this score and reduce randomness, we run 5 times of this test and the final score matrix was an average value of the 5 times replication (**Supplementary Figure S11b**). For example, in single needle case, the differentiation score between 10^-10^ M and 10^-12^ M is 0.7453, which means in 1000 rounds random comparison, among ∼745 times comparison, the binding number in 10^-10^ M larger than 10^-12^ M. These results mean that probability based binding events in Spectrum-CODE beads system (maximum binding number ∼11) have a theoretical ∼70% accuracy to successfully indicate the concentration status. Interestingly, when we further study the accuracy in binning condition (2×2 binning means we treat four rectangular layout needle as single ROI and the binding numbers will be summed together), we found the accuracy was significantly increased, especially higher than 4×4 binning, the accuracy was larger than 0.9999. This improvement effect of differentiation accuracy can be easily explained by Law of large numbers, where as the number of identically distributed, randomly generated variables increase, their sample mean (average) approaches their theoretical mean. The 4×4 binning including large number of binding particles (∼100-150 based on the experimental result) can evidently improve the analysis accuracy, which is also one of reasons that we choose 4×4 binning data for analysis even with the loss of a little spatial resolution.

#### What happened in multiplex condition when competition between different type of beads happened?

Here, we use 2 targets as an example and simulate the binding condition when two targets in different concentration compete (**Supplementary Figure S11c**). For starter, we compare the competition between two 10^-10^ M targets. Similar with the above simulation, two types of beads were assigned in the simulation area, where the binding probability were all set as 1/10. After 1000 times simulation, the binding number distribution on the needle surface of these two types of targets were summarized as a histogram. The distribution of these two targets (10^-10^ M vs 10^-10^ M) is almost similar. Further, we also simulated the condition where 10^-10^ M compete with 10^-12^ M, 10^-14^ M and 10^-16^ M. In these two types of targets with different concentration competition situation, our probability based binding events in spectrum-CODE beads system still presents identifiable binding number difference. The higher concentration targets specific beads mostly have higher binding number during the competition even in single needle. Combine with the above analysis, in a binning condition, this phenomenon will be more obvious.

### Note 4

#### miRNA and m^6^A-mRNA expression analysis among replication slices

We select three slice replications for miRNA profiling and m^6^A-mRNA profiling, including the dataset presented in the manuscript. Pearson correlation was calculated between every two pairs. The expression correlation of m^6^A-mRNA expression is much higher than that of miRNA (**Supplementary Figure S20**). Generally, miRNA can regulate gene networks dynamically through feedback and feedforward loops and steady state gene regulation. Besides, miRNA do not solely, or maybe even predominantly, function as target-specific regulators but may play key roles in the post-transcriptional reduction of expression noise^1^. The dynamic nature of miRNA expression might be the reason for relatively low correlation among bio samples, compared with that of mRNAs. Hypergeometric test was performed to calculate the enrichment significance in particular OB regions (**Supplementary Figure S21**). An expression feature fingerprint was generated for each cluster (For miRNA: 25×6 spatial expression feature vector for each cluster, where 1^st^ – 24^th^ rows are miRNA average expression in 6 brain region and 25^th^ row is the distribution ratio distribution in 6 brain region; For m^6^A-mRNA: 10×6 spatial expression feature vector for each cluster, where 1^st^ – 9^th^ rows are miRNA average expression in 6 brain region and 10^th^ row is the distribution ratio distribution in 6 brain region) and similarity between these clusters were calculated based on Euclidean distance of the corresponding expression feature fingerprint. All the cluster pairs with Euclidean distance <0.4 were plotted on the graph with edge connection. In miRNA cluster similarity network, we found cluster 3, 5 in manu batch, cluster 3, 6 in R1 batch and cluster 2 in R2 batch show inter connection with each other and interestingly, we found these clusters are all enriched in GL/OPL region. Representative spatial distribution of cluster 5 in manu batch, cluster 6 in R1 batch and cluster 2 in R2 batch are also presented with enrichment significance (-log10(p)). Similar GR enriched network and representative spatial distributions were also demonstrated in m^6^A-mRNA cluster similarity analysis (**Supplementary Figure S22**).

## Supplementary methods

### Immunofluorescent staining based single cell subtype clustering and mapping

The miRNAs or m^6^A mRNAs barcodes pattern was registered with the binarized astrocyte or neuron mask. The connected domains were analyzed with ‘bwlabel’ and ‘regionprops’ in Matlab. The copy number of the miRNAs or m^6^A mRNAs were calculated and normalized by dividing the correspond cell area. The single cell expression matrix was extracted and cluster analysis for the single cell subtype cluster was performed in R using the ConsensusClusterPlus package ^3^, where number of subsamples was set as 100, the proportion of items to sample was set as 0.8, the distance function was set as “euclidean” and k-means was used as the cluster algorithm. The optimized cluster number was selected based on the CDF delta area plot using the usual elbow method.

### Conjoint analysis of miRNA-mRNA spatial correlation and target relationship

Similar unsupervised cluster analysis was performed for both miRNAs and m^6^A mRNAs using BayesSpace R package^4^, where platform was set as “ST”, the mumber of top principal components to use when clustering was set as 7, the error model was set as “t”, the smoothing parameter was set as 2, the number of MCMC iterations was set as 10000. The enrichment of the clusters in related brain regions were calculated with hypergeometric test. Spatial distribution vector (SDV) of miRNAs or m^6^A mRNAs were calculated based on the distribution ratio of each cluster on the six brain regions. Then the correlation map of miRNAs and m^6^A mRNAs were calculated with the generated SDV for each two-cluster pair. Cluster pair with high correlation was selected for further analysis.

### Selection and design of m^6^A mRNA probes

Target mRNAs were designated using the database from Allen Brain Atlas ^5^. Top differential fold change mRNAs in main olfactory bulb with high confidence m6A sites were selected for the further profiling. m^6^A-Atlas ^6^ was used to identify the high-confidence m^6^A sites of specific mRNAs. Database from PA-m^6^A-seq, miCLIP, DART-seq, m^6^A-CLIP-seq, m^6^A-REF-seq, MAZTER-seq and m^6^A-seq with improved protocol were applied for reference. A 41 bp reference sequence was acquired and based on which, the related reporter probes for specific m^6^A mRNAs were designed.

### Spectral digitalization barcoding and encoding strategy

In our spectral digitalization coding system, fluorophores with different excitation wavelength or emission wavelength were selected for barcoding. 300-400 nm beads were used as coding media, which can be easily imaged in common confocal microscope. Machine learning (ML) algorithms were used for decoding with each fluorophore channel as a feature vector input. For starter, we considered a simple way to judge the existence of fluorophore directly by setting up a series of thresholds. However, the interference and crosstalk between adjacent channels makes this really hard. Sometimes, even we didn’t add the fluorophore, there will still be a peak influenced by other channels. Although using low crosstalk fluorophores may solve this, it will lead to a really small number of applicable fluorophore candidates. Employment of machine learning perfectly solved the crosstalk problem and make the application of adjacent fluorophore channels with strong crosstalk possible. ML decodes the fluorophore combinations by pattern recognition instead of isolated single threshold. Each barcode will have its specific spectrum pattern and can be decoded by machine learning algorithms.

Currently, 4 fluorophores were utilized with a total throughput of *N* × (2^C^− 1), where *C* is the number of fluorescent channels and N is the rounds of visualization cycles. With more fluorophores used, there will be exponential growth of the throughput. For example, with 7 fluorophore channels in single round, the throughput will be 127. Fluorophores with similar emission spectrum excited by different lasers or similar excitation lasers with different emission spectrum, can be used simultaneously in different imaging sequence, which increases the number of potential candidates. Here, we have listed seven fluorophore candidates with acceptable brightness available for a higher throughput probing.

Beside increasing more fluorophore channels, the mixed ratio of the fluorophores might also be adjusted. In this work, we use a binary mixing strategy where each channel only has two conditions, namely, ‘0’ or ‘1’. With larger mixed ratio number, for example, trinary mixing strategy, with ‘0’, ‘1’ and ‘2’, three conditions in each channel, there will surely be higher throughput. The equation will be *N* × (R_step_^C^ − 1), where R_step_ indicates the ratio step number, *C* indicates the fluorophore channels number, and *N* indicates the rounds of visualization cycles. In a trinary system with 7 different fluorophores would substantially increase the multiplexing throughput to over 10,000 by 5 rounds of visualization

### Machine learning (ML) based decoding algorisms

There were mainly two groups in the ML decoding process, namely the training group and the test group. When synthesizing a new batch of barcodes, parts of the barcodes with known labels will be treated as training group to extract the spectral features and training the ML model as well. The spectral feature extraction includes the following steps, namely capture coding signals, binarization, region of interest (ROI) selection and spectral pattern extraction. Here, in order to prepare the training group set, we use empty silicon biochip with glutaraldehyde functionalized to separately bind spectrum-CODE beads. Each type of coding has 3-5 biochips for ROI extraction and with more than 200 labelled ROIs were extracted for each spectrum-CODE. Finally, the training group signals and the test group signals all processed using extracted filter for background noise removal. In this case, the background signal noise with the similar shape and size of the particles will be cleaned. To extract an effective filter to reveal the barcodes spectral pattern, top 10% of the final extracted signals of the training group was used to calculate the corresponding filter. The upper boundary of the filter was calculated as

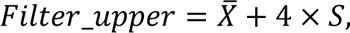

where 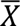 represented the average of the top 10% signals and *S* was the standard deviation of the top 10% signals. The lower boundary of the filter was calculated as

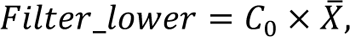

where *C*_0_ was a coefficient less than 1 and determined by the spectrum intensity range.

The ML was performed using classification toolbox in MATLAB, where ensembled bagged trees model was used with a learner type of decision tree. The maximum number of splits is 22087 and 30 learners were applied.

## Supplementary tables

**Table S1.**
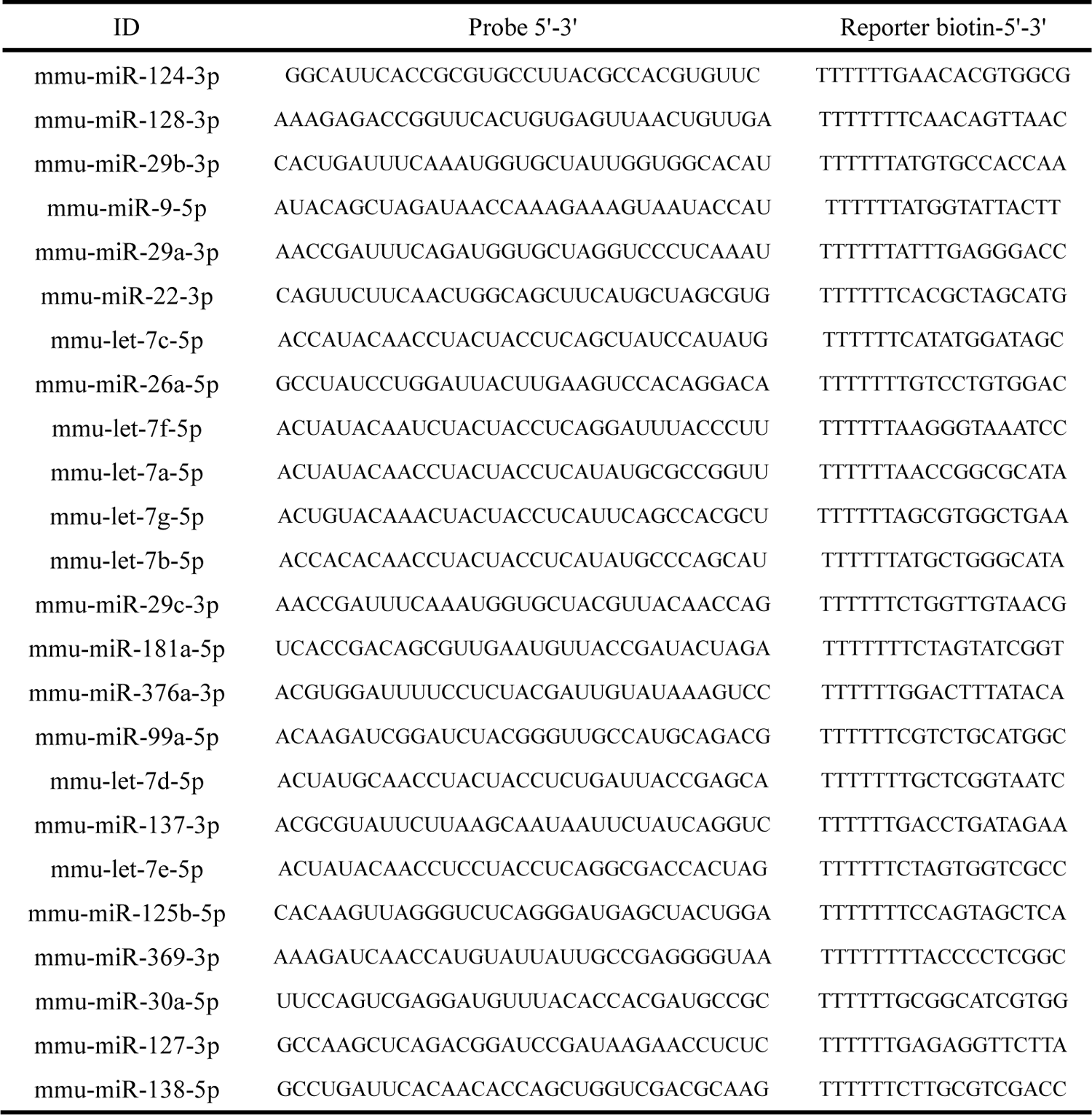
List examined miRNAs and their reporter sequences design.

**Table S2.**
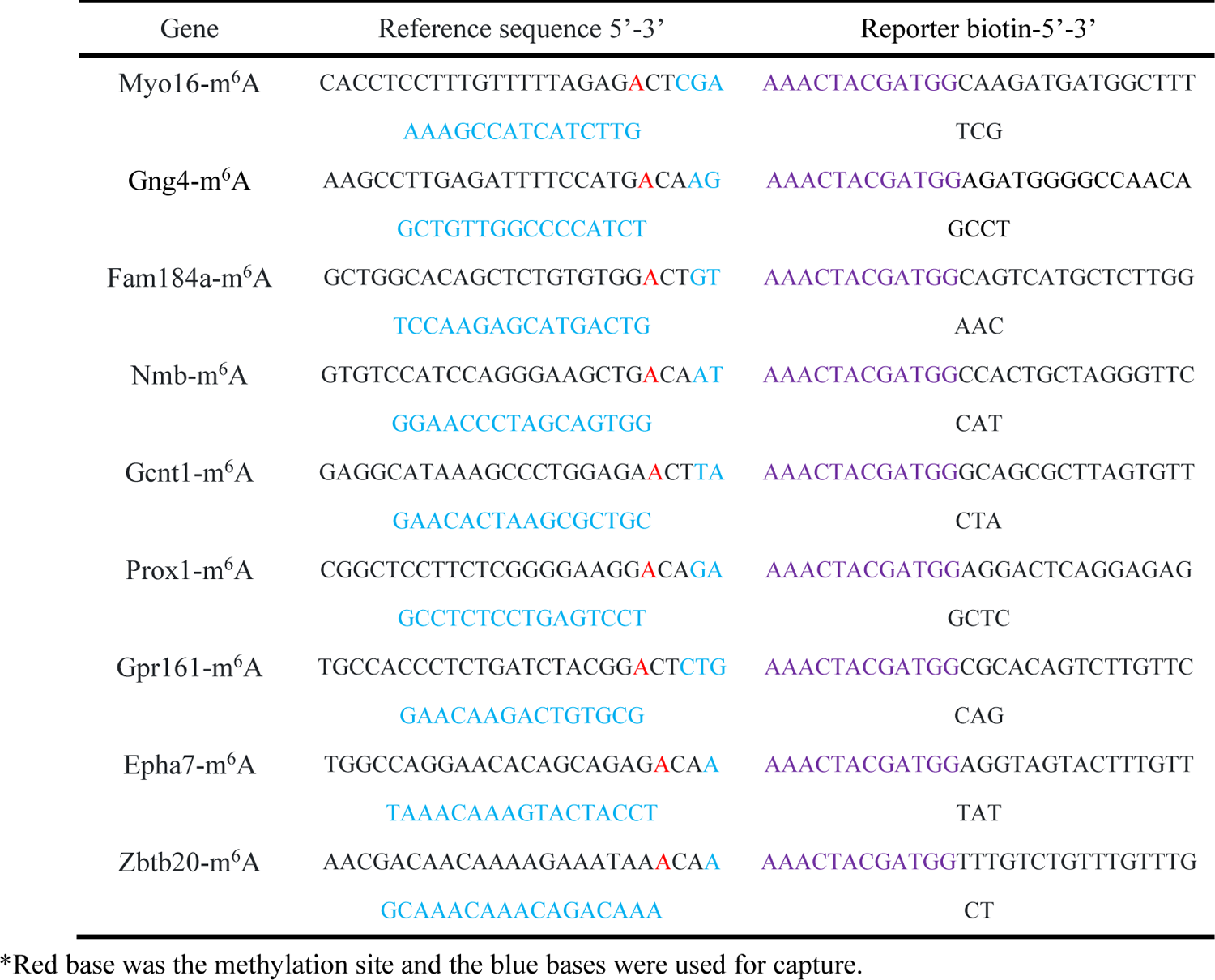
List examined m^6^A-mRNAs and their reporter sequences design.

